# A *Drosophila* larval premotor/motor neuron connectome generating two behaviors via distinct spatio-temporal muscle activity

**DOI:** 10.1101/617977

**Authors:** Aref Arzan Zarin, Brandon Mark, Albert Cardona, Ashok Litwin-Kumar, Chris Q. Doe

## Abstract

Animals generate diverse motor behaviors, yet how the same motor neurons generate distinct behaviors remains an open question. *Drosophila* larvae have multiple behaviors – e.g. forward crawling, backward crawling, self-righting and escape – and all of the body wall motor neurons (MNs) driving these behaviors have been identified. Despite impressive progress in mapping larval motor circuits, the role of most motor neurons in locomotion remains untested, the majority of premotor neurons (PMNs) remain to be identified, and a full understanding of proprioceptor-PMN-MN connectivity is missing. Here we report a comprehensive larval proprioceptor-PMN-MN connectome; describe individual muscle/MN phase activity during both forward and backward locomotor behaviors; identify PMN-MN connectivity motifs that could generate muscle activity phase relationships, plus selected experimental validation; identify proprioceptor-PMN connectivity that provides an anatomical explanation for the role of proprioception in promoting locomotor velocity; and identify a new candidate escape motor circuit. Finally, we generate a recurrent network model that produces the observed sequence of motor activity, showing that the identified pool of premotor neurons is sufficient to generate two distinct larval behaviors. We conclude that different locomotor behaviors can be generated by a specific group of premotor neurons generating behavior-specific motor rhythms.

## Introduction

Locomotion is a rhythmic and flexible motor behavior that enables animals to explore and interact with their environment. Birds and insects fly, fish swim, limbed animals walk and run, and soft-body invertebrates crawl. In all cases, locomotion results from coordinated activity of muscles with different biomechanical output. This precisely regulated task is mediated by neural circuits composed of motor neurons (MNs), premotor interneurons (PMNs), proprioceptors, and descending command-like neurons (Marder and Bucher 2001; Arber 2017; Arber and Costa 2018). A partial map of neurons and circuits regulating rhythmic locomotion have been made in mouse (Crone et al. 2008; Grillner and Jessell 2009; Zagoraiou et al. 2009; Dougherty et al. 2013; Goetz et al. 2015; Bikoff et al. 2016), cat (Kiehn 2006; Nishimaru and Kakizaki 2009), fish (Kimura et al. 2013; Song et al. 2016), tadpole (Roberts et al. 2008; Roberts et al. 2010), lamprey (Grillner 2003; Mullins et al. 2011), leech (Brodfuehrer and Thorogood 2001; Kristan et al. 2005; Marin-Burgin et al. 2008; Mullins et al. 2011), crayfish (Mulloney and Smarandache-Wellmann 2012; Mulloney et al. 2014), and worm (Tsalik and Hobert 2003; Wakabayashi et al. 2004; Haspel et al. 2010; Kawano et al. 2011; Piggott et al. 2011; Wen et al. 2012b; Zhen and Samuel 2015; Roberts et al. 2016). These pioneering studies have provided a wealth of information on motor circuits, but with the exception of *C. elegans* (White et al. 1986), there has been no system where all MNs and PMNs have been identified and characterized. Thus, we are missing a comprehensive picture of how an ensemble of interconnected neurons generate diverse locomotor behaviors.

We are interested in understanding how the *Drosophila* larva executes multiple behaviors, in particular forward versus backward locomotion (Carreira-Rosario et al. 2018). Are there different motor neurons used in each behavior? Are the same motor neurons used but with distinct patterns of activity determined by premotor inputs? How does the ensemble of premotor and motor neurons generate additional behaviors, such as escape behavior via lateral rolling? A rigorous answer to these questions requires both comprehensive anatomical information – i.e. a premotor/motor neuron connectome – and the ability to measure rhythmic neuronal activity and perform functional experiments. All of these tools are currently available in *Drosophila*, and we use them to characterize the neuronal circuitry used to generate forward and backward locomotion, and how proprioception is integrated by the PMN ensemble.

The *Drosophila* larva is composed of 3 thoracic (T1-T3) and 9 abdominal segments (A1-A9), with sensory neurons extending from the periphery into the CNS, and motor neurons extending out of the CNS to innervate body wall muscles (Figure 1A). Most segments contain 30 bilateral body wall muscles that we group by spatial location and orientation: dorsal longitudinal (DL; includes previously described DA and some DO muscles), dorsal oblique (DO), ventral longitudinal (VL), ventral oblique (VO), ventral acute (VA) and lateral transverse (LT)(Figure 1B)(Crossley 1978; Hooper 1986; Bate 1990). Using these muscles, the larval nervous system can generate forward locomotion, backward locomotion, turning, hunching, digging, self-righting, and escape (reviewed in Kohsaka et al. 2017; Clark et al. 2018). Here we focus on forward and backward locomotion. Forward crawling behavior in larvae involves a peristaltic contraction wave from posterior to anterior segments; backward crawling entails a posterior propagation of the contraction wave (Crisp et al. 2008; Dixit et al. 2008; Berni et al. 2012; Gjorgjieva et al. 2013; Heckscher et al. 2015; Pulver et al. 2015; Loveless et al. 2018).

**Figure 1.**
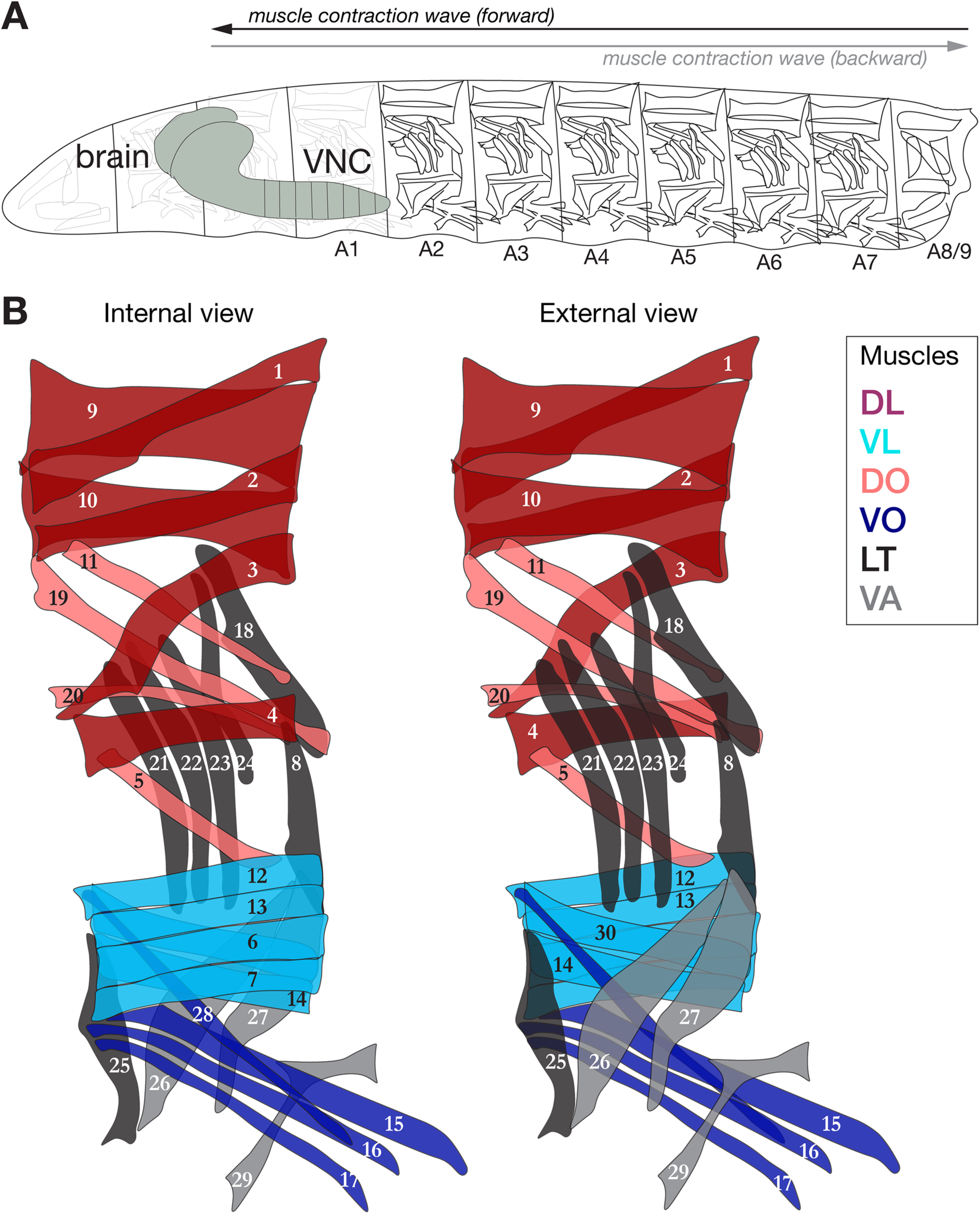
Schematic depiction of the larval neuromuscular system. (A) *Drosophila* larva contain three thoracic and nine abdominal segments, the muscles of which are innervated by MNs located in the corresponding thoracic and abdominal segments. (B) Schematic of the 30 muscles of abdominal segments (A2-A6) from internal and external view. Segment A1 is similar to A2-A6, with the exception that it lacks muscle 25 and MN-25.

Body wall muscles are innervated by approximately 60 MNs per segment, consisting of 28 left/right pairs that typically each innervate one muscle, and whose neuromuscular junctions have <u>b</u>ig boutons, therefore also called type-I<u>b</u> MNs; two pair of type-Is (<u>s</u>mall bouton) MNs that innervate large groups of dorsal or ventral muscles; three type II ventral unpaired median MNs that provide octopaminergic innervation to most muscles; and one or two type III insulinergic MNs innervating muscle 12 (Gorczyca et al. 1993; Landgraf et al. 1997; Hoang and Chiba 2001; Landgraf et al. 2003; Choi et al. 2004; Mauss et al. 2009; Koon et al. 2011; Koon and Budnik 2012; Zarin and Labrador 2017). All MNs in segment A1 have been identified by backfills from their target muscles (Landgraf et al. 1997; Landgraf et al. 2003; Mauss et al. 2009), and several have been shown to be rhythmically active during larval locomotion, but only a few of their premotor inputs have been described (Kohsaka et al. 2014; Heckscher et al. 2015; Fushiki et al. 2016; Hasegawa et al. 2016; Zwart et al. 2016; Takagi et al. 2017; Carreira-Rosario et al. 2018). Some excitatory PMNs are involved in initiating activity in their target MNs (Fushiki et al. 2016; Hasegawa et al. 2016; Zwart et al. 2016; Takagi et al. 2017; Carreira-Rosario et al. 2018), while some inhibitory PMNs limit the duration of MN activity (Kohsaka et al. 2014; MacNamee et al. 2016; Schneider-Mizell et al. 2016) or produce intrasegmental activity offsets (Zwart et al. 2016). Interestingly, some PMNs are active specifically during forward locomotion or backward locomotion (Kohsaka et al. 2014; Heckscher et al. 2015; Fushiki et al. 2016; Hasegawa et al. 2016; Takagi et al. 2017; Carreira-Rosario et al. 2018). In addition, there are six pair of proprioceptor neurons in each abdominal segment (ddaE, ddaD, vpda, dmd1, dbd and vbd). They are important for promoting locomotor velocity and posture (Hughes and Thomas 2007; Heckscher et al. 2015; Ohyama et al. 2015; Burgos et al. 2018; He et al. 2019; Vaadia et al. 2019), and some of their CNS targets have been identified (Heckscher et al. 2015; Ohyama et al. 2015), but to date little is known about how or if they are directly connected to the PMN/MN circuits.

## Results

### TEM reconstruction of all segmental motor neurons

Understanding how the CNS generates distinct behaviors using the same neural circuitry is an important question in neuroscience. One of the first steps in answering this question is to define the neural circuits used to generate the relevant behaviors, ideally with single synapse resolution as a comprehensive connectome. A connectome can provide testable hypothesis, and constrain theoretical models (Bargmann and Marder 2013). Recently, a serial section transmission electron microscopy (TEM) volume of the entire first instar larval CNS was generated (Ohyama et al. 2015). To date, however, only a small fraction of PMNs and MNs have been reconstructed (Heckscher et al. 2015; Fushiki et al. 2016; Zwart et al. 2016; Carreira-Rosario et al. 2018). Here, we identify and reconstruct all differentiated MNs in segment A1, which can be used as a proxy for other abdominal segments. We identified all 26 pair of type Ib MNs with single or dual muscle targets, the three unpaired midline octopaminergic MNs, both pair of type Is MNs that target large muscle groups (RP2, RP5), and one pair of type III MN (Figure 2; Table 1). The presence of yet another type Is MN has been suggested (Hoang and Chiba 2001), but we did not find it in the TEM volume; it may be late-differentiating or absent in A1. We linked all 31 bilateral MNs in the TEM volume to their muscle target by matching the dendritic morphology in the EM reconstruction to the dendritic morphology determined experimentally (Landgraf et al. 1997; Landgraf et al. 2003; Mauss et al. 2009) (Figure 2; Figure 2 – figure supplements 1, 2; Table 1). A dataset of all MNs that can be opened in CATMAID (Saalfeld et al. 2009) is provided as Supplemental File 1. Note that the transverse nerve MN (MN25-1b) is only present in the A2-A7 segments (Hessinger et al. 2017), so we traced it in A2. We found that all MNs had a dense array of post-synapses on their dendritic projections, but unlike *C. elegans* (Wen et al. 2012b) we observed no pre-synaptic contacts to other MNs or interneurons (Figure 2 – figure supplements 1, 2). In conclusion, we have successfully identified and reconstructed, at single synapse-level resolution, all differentiated MNs in segment A1 of the newly hatched larval CNS.

**Table 1.**
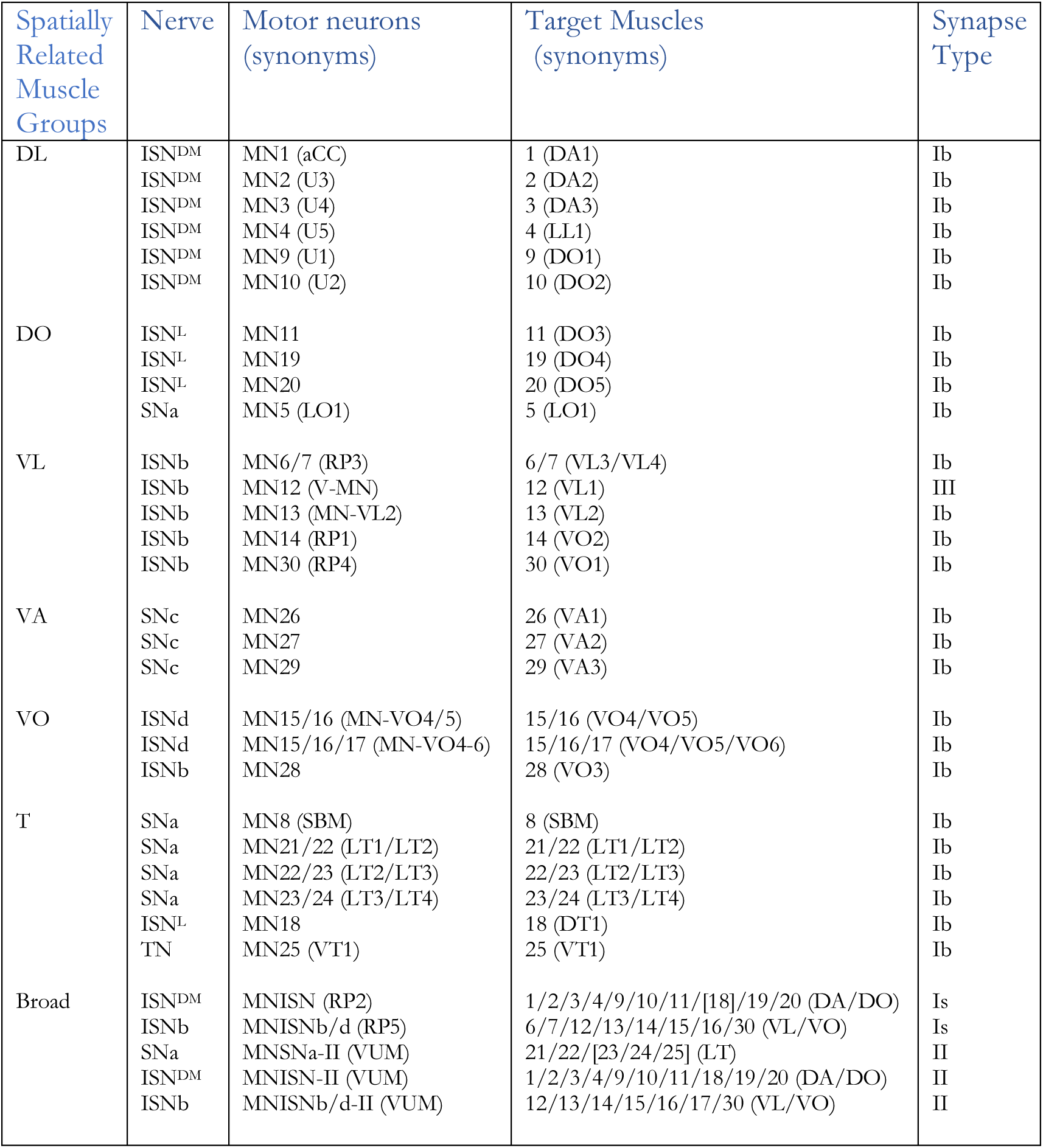
Motor neurons present in the CATMAID reconstruction. All MNs were identified in the first abdominal segment on both left and right sides, with the exception of MN25 which is not present in A1 and thus annotated in A2. See text for abbreviations.

**Figure 2.**
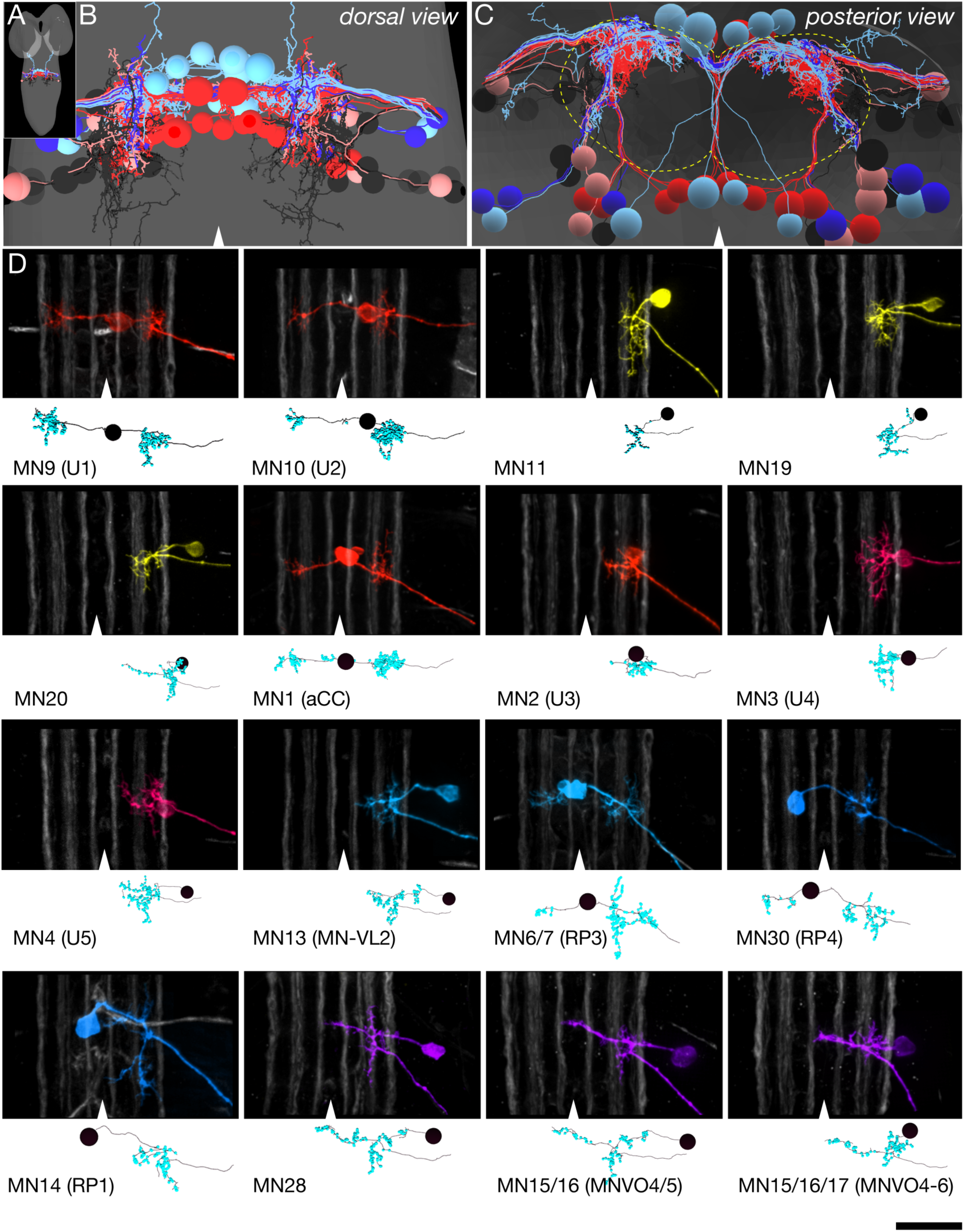
Identification of all differentiated motor neurons in segment A1 of the TEM volume. (A) Dorsal view of the TEM reconstruction of the L1 CNS (gray shading) showing all bilateral MNs reconstructed at single synapse level. The one intersegmental dendrite is from RP3 in A1; it is not observed in other abdominal segments. (B) Dorsal view of centered on the A1 segment; midline, arrowhead. MNs are color-coded as in Figure 1B. (C) Posterior (cross-section) view of the neuropil (outlined) and cortex in A1. Note the MN dendrites target the dorsal neuropil. (D) Representatives showing the morphological similarity between MNs identified in vivo by backfills (Mauss et al. 2009) versus the most similar MN reconstruction from the TEM volume. The top section in each panel shows the morphology of the MN dendrites based on in vivo backfills; used with permission); six distinct Fas2 fascicles (three per hemisegment) are shown in white; midline, arrowhead. The bottom section shows MN dendrite morphology reconstructed from the TEM volume in A1.

### TEM reconstruction of 67 premotor neurons

We next identified the tier of PMNs with monosynaptic contacts to MNs in segment A1. This included local premotor neurons with somata in A1 as well as neurons from adjacent segments with dense connectivity to A1 MN dendrites. We identified 67 bilateral PMNs (134 total) with connectivity to A1 MNs (Figure 3A,B; Table 2; see Methods for selection criteria). The PMN cell bodies were distributed throughout the segment, and sent projections into the dorsal neuropil (Figure 3A,B; Figure 3 – supplement 1), with pre-synaptic sites strongly enriched in the dorsal neuropil (Figure 3C,D; Figure 3 – supplement 1). This is expected, as the motor neuron dendrites target the dorsal neuropil (Landgraf et al. 1997; Mauss et al. 2009).

**Table 2.**
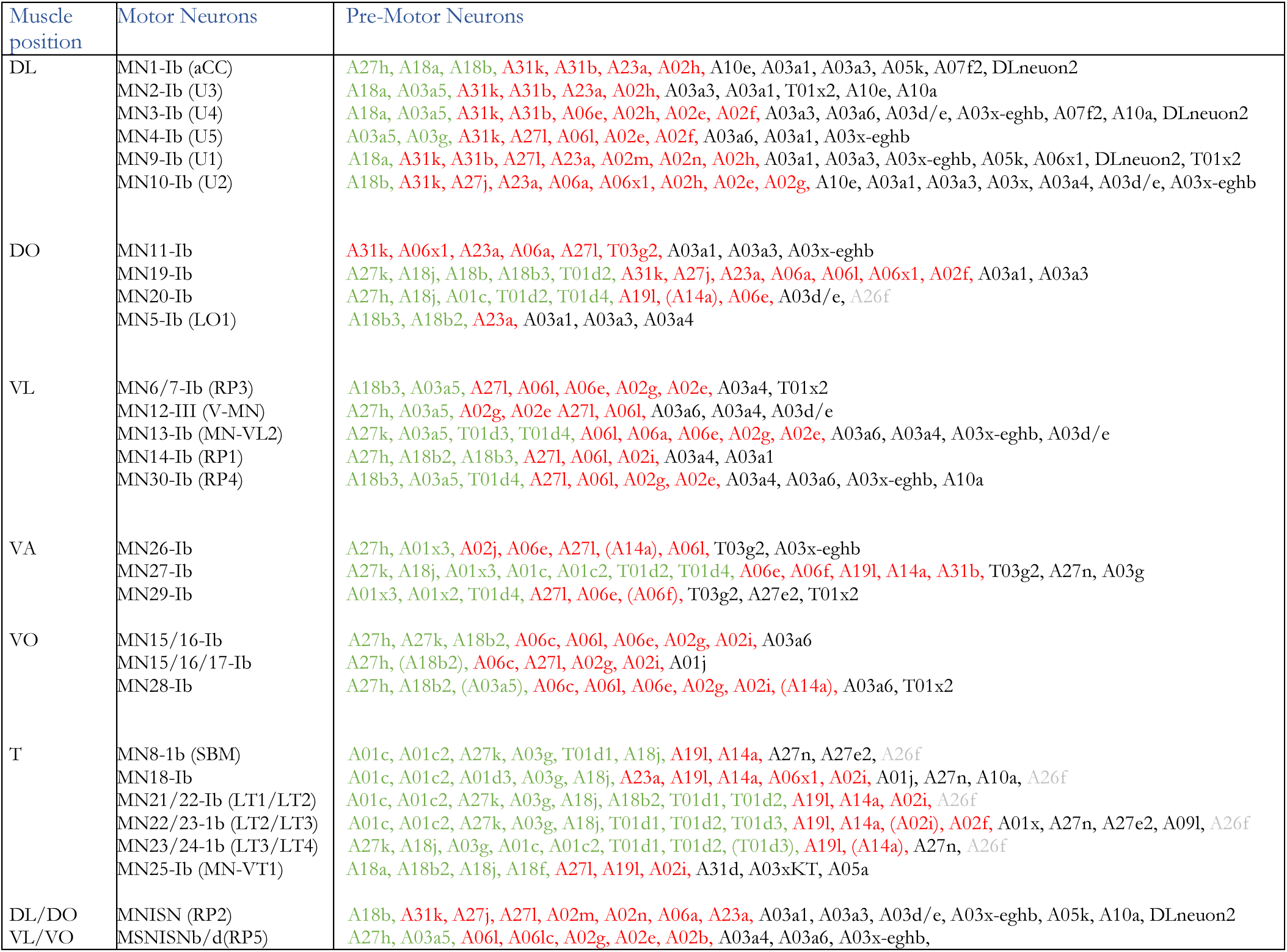
Premotor neurons innervating type Ib MNs. Left column, SMuGs. Middle column, type Ib MNs innervating 1-3 muscles in each muscle group (synonym, parentheses); the widely innervating type Is MNs RP2 and RP5 are not shown. Right column, premotor interneurons innervating the indicated MNs (green, presumed excitatory; red, presumed inhibitory; grey, corozonergic; black, unknown. Premotor connectivity uncertain, parentheses.

**Figure 3.**
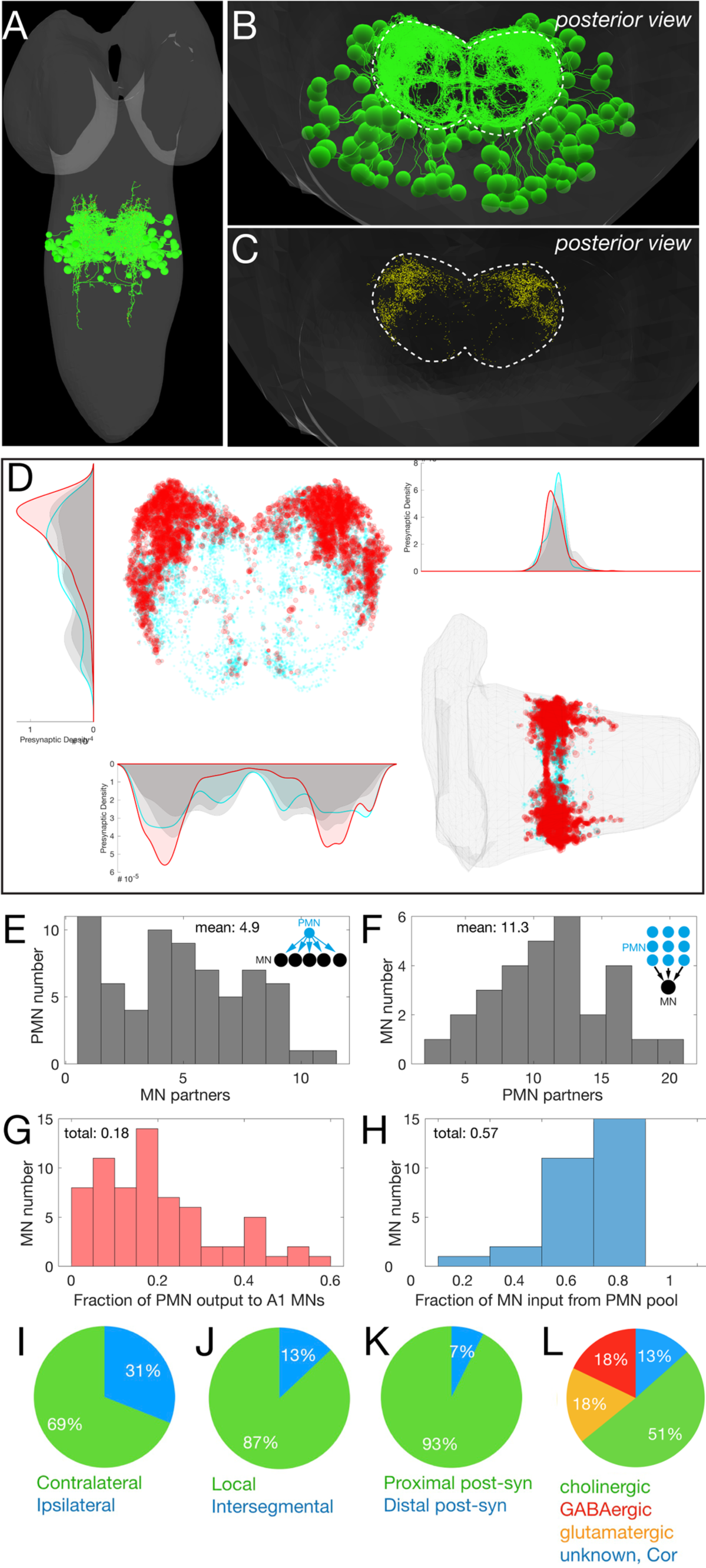
Identification of 67 premotor neurons at synapse-level in the EM reconstruction. (A) Dorsal view of centered on the A1 segment showing all 67 pair of PMNs reconstructed in this study. (B) Posterior (cross-section) view of the neuropil (outlined) and cortex in A1. Note the PMN cell bodies are located in the cortex while their dendrites target the dorsal neuropil. Dorsal, up; midline, arrowhead. (C) Posterior (cross-section) view of the neuropil (outlined) and cortex in A1. Every PMN pre-synaptic site is labeld (yellow dots). Note how the PMN pre-synapses are enriched in the dorsal neuropil. Dorsal, up. (D) Spatial distributions of pre- and post-synaptic sites for all PMNs. Plots are 1D kernel density estimates. Each red dot represents a single pre-synaptic site scaled by its number of outputs. Each cyan dot represents a single post-synaptic site. Left: posterior view, dorsal up. Right: dorsal view; anterior left. (E-H) Quantification of PMN-MN connectivity. (E) PMNs innervate an average of 4.9 A1 MNs. (F) MNs receive inputs from an average of 11.3 PMNs from this population of PMNs. (G) Histogram showing 18% of PMN output onto MNs. H) Histogram showing 57% of MN post-synapses receive input from the 67 PMNs, with a range from <20% to nearly 80%. (I-L) Quantification of PMN morphology and neurotransmitter expression.

We observed widespread connectivity of PMNs to multiple MNs (Figure 3E). Each PMN synapsed with an average of 4.9 MNs (Figure 3E), and each MN had an average of 11.3 input PMNs (Figure 3F). All PMNs targeted both MNs and interneurons; there were no PMNs exclusively innervating MNs (Figure 3G). The 67 bilateral PMNs make 5599 synapses associated with A1 MNs which account for 18% of PMN output and 57% of the total A1 MN input (Figure 3H). In addition, most PMNs projected contralaterally, had local arbors, and had post-synaptic inputs on their more proximal processes (Figure 3I-K). The few PMNs with pre- and post-synapses clustered distally (Figure 3 – supplement 1C) are good candidates for nonspiking interneurons that perform local computations (reviewed in Pearson 1976; Marder and Bucher 2001). A file that can be opened in CATMAID showing all 67 bilateral PMNs is provided as Supplemental File 2. Thus, we have identified the majority of the PMN inputs to the A1 MN population.

Neurotransmitter expression has been characterized for only a small fraction of the PMNs described here (Kohsaka et al. 2014; Heckscher et al. 2015; Fushiki et al. 2016; Hasegawa et al. 2016; MacNamee et al. 2016; Zwart et al. 2016; Takagi et al. 2017; Yoshino et al. 2017; Burgos et al. 2018; Carreira-Rosario et al. 2018). To increase coverage, we screened for Gal4 lines with sparse expression patterns, performed MultiColorFlpOut (Nern et al. 2015) to match their morphology to individual PMNs, and mapped neurotransmitter expression. We found 12 GABAergic (presumptive inhibitory), 9 glutamatergic (presumptive inhibitory), 34 cholinergic (presumptive excitatory), and 1 corozonergic (neuromodulatory) neurons; 11 PMNs could not be identified due to lack of Gal4 lines (Figure 3L; Supplemental Table 2), and we did not identify any neurons co-expressing two fast neurotransmitters.

Thus, we have identified 67 bilateral PMNs which innervate the MNs in segment A1, mapped all PMN-MN synapses within segment A1, and determined neurotransmitter expression for the majority of the PMNs. We conclude that PMNs target the majority of their pre-synapses to the dorsal neuropil, where they connect to other interneurons, PMNs, and MNs.

### All body wall muscles are activated during forward and backward locomotion

The PMN-MN connectome supports diverse motor behaviors, including forward and backward locomotion. To understand how these PMN-MN circuits generate different behaviors requires mapping neuronal activity to establish a neuron-behavior map. To map MN activity, we took advantage of the fact that each of the type Ib MNs typically innervates a single muscle, and thus muscle depolarization can be used as a proxy for the activity of its innervating MN. To date only muscle contraction, rather than muscle activity, data have been collected, and only for five of the 30 body wall muscles (Heckscher et al. 2012; Zwart et al. 2016). Muscle contraction could occur passively due to biomechanical linkage between adjacent muscles, so it is not a perfect substitute for directly measuring muscle activity. Thus, it remains unknown whether some or all muscles are activated by MNs during locomotion, nor has the phase relationship between most muscles been determined. Both are important for understanding how rhythmic motor patterns are generated in this system.

We used GCaMP6f/mCherry ratiometric calcium imaging to measure the activation time of all 30 individual body wall muscles in the abdominal segments. We expressed GCaMP6f and mCherry using the muscle line *R44H10-LexA*, which has variable expression in sparse to dense patterns of muscles. For this experiment we analyzed larvae with dense muscle expression. We imaged both forward and backward locomotion in 2nd instar larvae (a representative animal shown in Figure 4A, D). We found that an increased GCaMP6f signal correlated with muscle contraction during both forward and backward locomotion (representative examples of muscle 6 shown in Figure 4B, E). Most importantly, all imaged muscles (30 for forward and 29 for backward) showed a significant rise in GCaMP6f fluorescence during forward and backward locomotion (Figure 4C, F; Movies 1, 2). We conclude that all MNs and their target muscles are activated during forward and backward locomotion.

**Figure 4.**
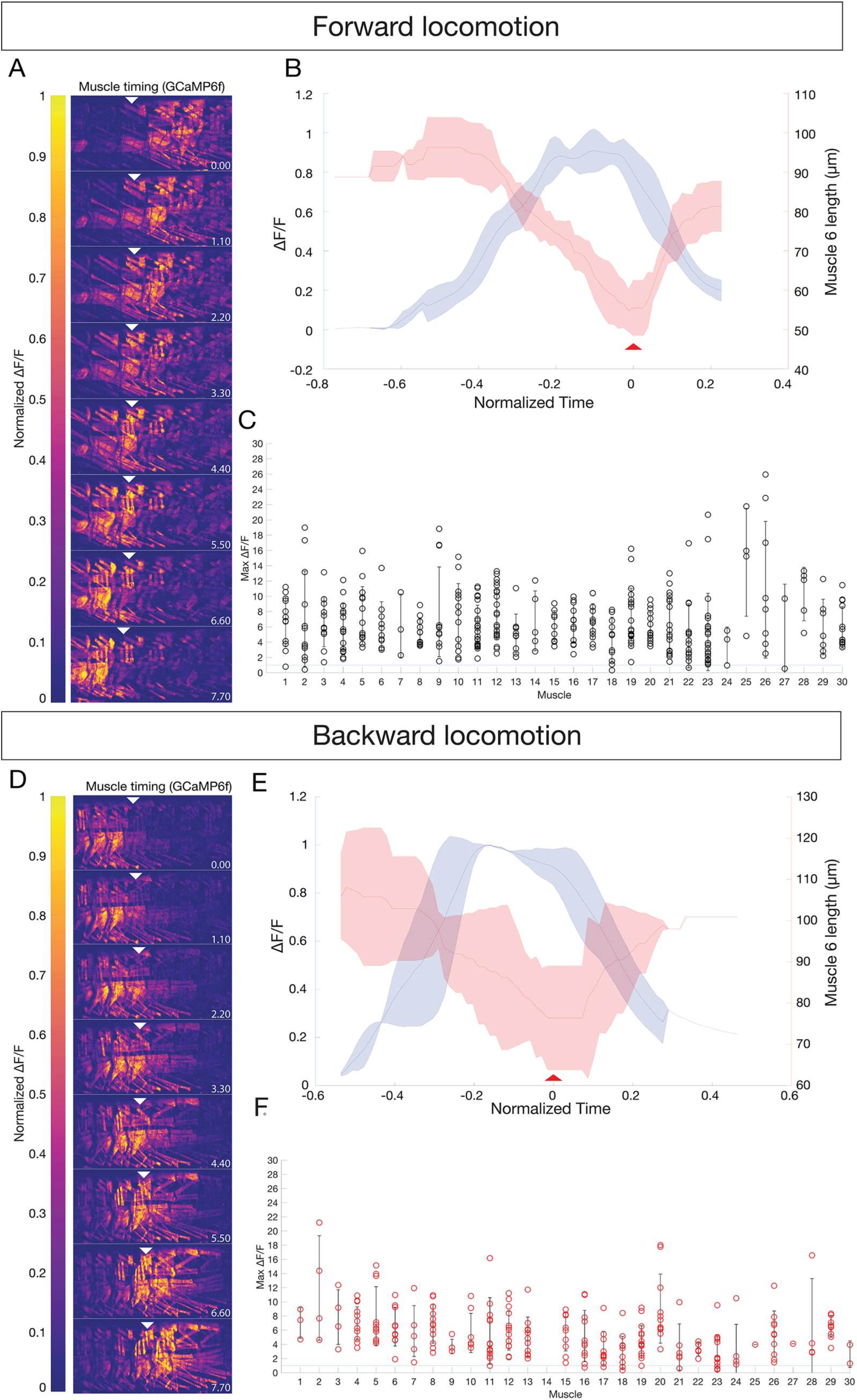
All body wall muscles are utilized during forward and backward locomotion. (A,D) Sequential images of muscle GCaMP6f ΔF/F signal during forward (A) or backward (D) locomotion. GCaMP6f levels were normalized to mCherry. Anterior to left, dorsal up; time in seconds. Genotype: *GMR44H10-LexA lexAOP-GCaMP6f; -LexA lexAOP--mCherry.* (B,E) Mean calcium transient (blue) vs mean muscle length (red) measurements for muscle 6 during forward (B) or backward (E) locomotion. N = 3 segments. T_0_ was set as the point of maximum contraction as determined by muscle length for each crawl. Shaded bars represent standard deviation. (C,F) All observed muscles show calcium transients greater than 100% ΔF/F during forward (C) or backward (F) locomotion. Each dot represents the maximum GCaMP ΔF/F signal in the indicated muscle during a single crawl, normalized to mCherry. Error bars represent standard deviation. Muscle names as in Figure 1.

### Hierarchical clustering identifies MN phase relationships during forward and backward locomotion

To determine the timing of individual MN/muscle activation during forward and backward locomotion, we embedded the multidimensional crawl cycle data in two-dimensional space using principal component analysis (PCA)(Lemon et al. 2015). We aligned crawl trials by finding peaks in this 2D space which corresponded to the highest amplitude of the most muscles in a given crawl (Figure 5 – figure supplement 1; see Methods). Although muscle activity appeared as a continuum with the sequential recruitment of individual muscles (Figure 5A), hierarchical clustering of the mean activity of each muscle during forward and backward crawling revealed four groups of co-active muscles for both behaviors (Figure 5B-E; summarized in Figure 5F,G; Table 3). We call these <u>C</u>o-active <u>Mu</u>scle <u>G</u>roups or CMuGs. Analysis of forward locomotion showed that each CMuG had different patterns of activation: e.g. CMuG F1 had a more variable time of onset, whereas CMuG F4 had a highly coherent onset (Figure D-E). Furthermore, the time of CMuG activation was more coherent than the time of its inactivation (Figure 5D-G). Overall, longitudinal muscles tended to be active early in the crawl cycle, and transverse muscles activated late in the crawl cycle (Figure 5A-E), consistent with prior reports tracking single muscles within each group (Heckscher et al. 2012; Zwart et al. 2016). There were exceptions, however. Several ventral longitudinal muscles and dorsal oblique muscles, presumably with different biomechanical functions due to their different orientation and position, were synchronously recruited in CMuG F2 (Figure 5F; Table 3).

**Table 3.** Co-activated muscle groups during forward or backward locomotion. There are four co-activated muscle groups during backward and forward locomotion, but the muscles in each group differ in forward versus backward locomotion. Note that backward locomotion is not simple a reverse of the pattern seen in forward locomotion. This represents the most common activation sequences, although there is some variation, particularly during the fastest locomotor velocities.

| <u>Forward</u> | <u>Co-activated muscles</u> |
| --- | --- |
| F1 | 2,6,10,11,14,30 |
| F2 | 3,4,5,9,12,13,18,19,25,26,29 |
| F3 | 1,8,15,16,17,20,28 |
| F4 | 21,22,23 |
| <u>Backward</u> | <u>Co- activated muscles</u> |
| B1 | 10,15,16,17 |
| B2 | 1,3,4,6,9,12,13,28 |
| B3 | 2,5,8,19,20,26,29 |
| B4 | 11,18,21,22,23,24 |

**Figure 5.**
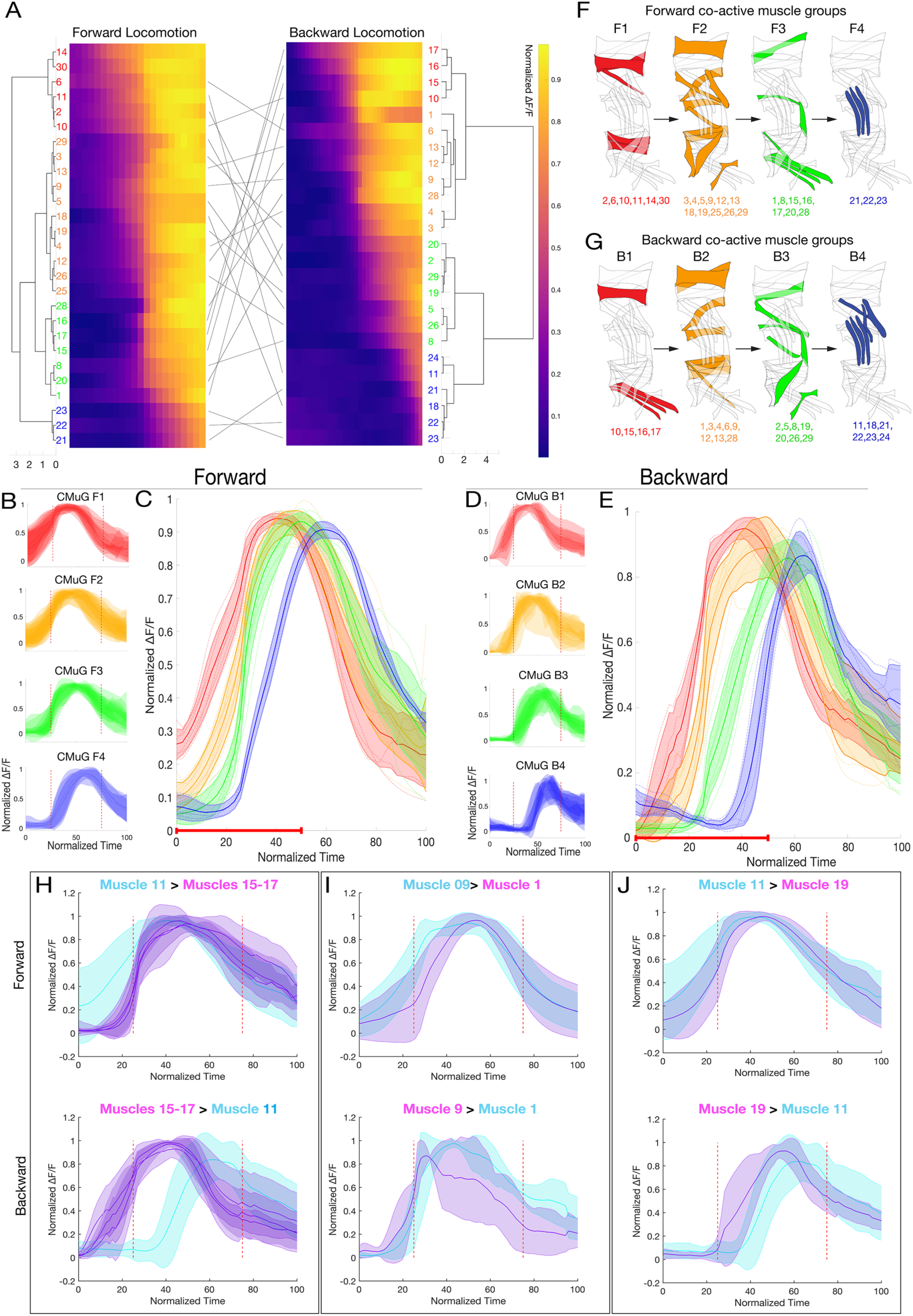
Larval body wall muscles form four co-activated muscle groups during forward and backward locomotion. (A) Hierarchical clustering of mean activity for all observed muscles yields four CMuGs during forward locomotion (F1-F4) and a different group of four during backward locomotion (B1-B4). Heatmaps represent the mean range-normalized calcium activity of each muscle (n > 3 forward crawl bouts for each muscle, with a total of 337 individual muscles analyzed across 23 crawls for forward and 188 individual muscles analyzed across 14 crawls for backward locomotion). Clustering was performed only on the first half of the crawl cycle to maximize the contribution of muscle onset time in determining CMuG membership. Cluster number was determined by visual inspection of the dendrogram as well as the gap-criterion optimal cluster number. Black lines are a tanglegram of individual muscle locations in the crawl cycle. (B) Plots of average muscle activity for muscles in each forward CMuG. Error bars represent the standard deviation of individual muscles. (C) Plots of average forward CMuG activity timing. Error bars represent the standard deviation of the average muscle activity of each muscle in a given CMuG. Dotted lines represent the average muscle activity for each muscle in a given CMuG. Red line along the x-axis represents the fraction of the crawl cycle that was used for clustering. (D) Plots of average muscle activity for muscles in each backward CMuG. Error bars represent the standard deviation of individual muscles. (E) Plots of average backward CMuG activity timing. Error bars represent the standard deviation of the average muscle activity of each muscle in a given CMuG. Dotted lines represent the average muscle activity for each muscle in a given CMuG. Red line along the x-axis represents the fraction of the crawl cycle that was used for clustering. (F) Schematic representation of the CMuGs for forward locomotion. (G) Schematic representation of the CMuGs for backward locomotion. (H) During forward locomotion, muscle 11 is activated before muscle 15-17, while their order is flipped during backward crawling. (I) During forward locomotion, muscle 1 is active after muscle 9, whole they become synchronously active during backward locomotion. (J) During forward locomotion, muscle 11 is activated before muscle 19, while their order is flipped during backward crawling.

Importantly, CMuGs are not the same as the previously described spatially-related muscle groups, which we call <u>S</u>patial <u>Mu</u>scle <u>G</u>roups or SMuGs (see Figure 1). The muscles in each SMuG are thought to provide a similar biomechanical function, due to the their similar position and orientation along the body wall, and published data on several individual muscles was consistent with each SMuG having a similar time of activity (Heckscher et al. 2015; Zwart et al. 2016). However, our results show that SMuGs can be asynchronously recruited: dorsal longitudinal muscles, presumably with similar biomechanical functions, are part of three different CMuGs (F1-F3) (Figure 5B; Table 3). This is surprising, as we expected that muscles with similar position and orientation – and thus presumably with similar functions – to be co-activated. Our finding that co-active muscle groups (CMuGs) are distinct from SMuGs raises several questions we address below: Do MNs that innervate each CMuG localize their dendrites to a discrete region of the neuropil, as reported for SMuGs (Landgraf et al. 2003; Mauss et al. 2009)? Are there distinct pools of PMNs for each SMuG and CMuG? Do proprioceptors provide differential input to SMuGs or CMuGs?

We performed the same analysis for backward locomotion, to determine if it was simply the reverse of forward or a completely different pattern (and behavior). During backward locomotion, muscle activity also clusters into four CMuGs, but the muscles in each group are different than for forward locomotion, so we call them B1-B4 (Figure 5A,D,E,G; Table 3). Generally, longitudinal muscles tended to be active early in the crawl cycle, and transverse muscles activated late in the crawl cycle, during both forward and backward locomotion (Figure 5F,G). Plotting the average time of muscle activity showed that muscles in CMuG B3 and B4 had a highly coherent rise of activity, whereas CMuGs B1 and B2 were more variable (Figure 5D,E). As with forward locomotion, muscles in different SMuGs can be synchronously recruited (e.g. the DL and VO muscles 10, 15-17 in CMuG B1)(Figure 5G; Table 3). Conversely, muscles in the same SMuGs can be asynchronously recruited (e.g. the DL muscles 1, 2, 9, 10, are found in CMuGs B1-B3 (Figure 5G; Table 3). Overall, we find that the pattern of muscle activation during backward crawling is distinct from that of forward crawling (Figure 5F,G). However, there are also some similarities. For example, muscles 21-23 are in the last CMuG and muscle 10 is in the first CMuG, during both forward and backward locomotion (Figure 5F,G). A comparison of recruitment times for all muscles imaged in both forward and backward locomotion is provided in Figure 5 – figure supplement 2. We conclude that forward and backward locomotion are two distinct motor patterns, not simply the same pattern in reverse, and that pre-motor/motor circuitry has the ability to drive two distinct patterns of rhythmic muscle activity during the two different behaviors. Similar to forward locomotion, CMuGs during backward locomotion do not match SMuGs.

### Motor neurons can generate CMuGs without clustering of post-synaptic inputs

Next, we wanted to understand the underlying mechanisms for how CMuGs become temporally segregated during locomotion. One possibility is that MNs in a specific CMuG target their post-synaptic sites to a common neuropil domain. To address this question, we clustered MNs based on the similarity of the spatial distributions of their inputs (Schlegel et al. 2016). We first compared MNs in the left and right A1 hemisegments and observed highly similar post-synapse clustering (Pearson correlation coefficient, *r* = 0.97), which we averaged for subsequent analysis. This validated the quality and reproducibility of the MN dendritic reconstructions and highlighted the stereotypy of MN synaptic locations in the neuropil. We mapped the locations of the inputs that MNs receive within the neuropil, comparing MNs innervating different SMuGs (Figure 6A,B) or different CMuGs (Figure 6C,D). Consistent with previously published single muscle data (Heckscher et al., 2015; Zwart et al., 2016), we found significantly different localization of post-synaptic input sites for MNs innervating the spatial muscle groups DL, DO, VL, and VO compared to LT and VA muscle groups (Figure 6A), Similarly, different post-synaptic input sites were seen for MNs innervating VA compared to LT muscles (Figure 6B). These results confirm and extend previous reports of MN myotopic maps (Landgraf et al. 2003; Mauss et al. 2009), but now at an individual synapse level of resolution. We also observed segregation in post-synapse positions between F4 and other forward CMuGs (F1, F2, F3) as well as between B4 and other backward CMuGs (B1, B2, B3) (Figure 6C-D), but this is likely due to the F4 and B4 CMuGs overlapping considerably with the LT muscles, which are known to be active at the end of each crawl cycle (see above; Heckscher et al., 2015; Zwart et al., 2016). Otherwise, we saw no obvious segregation of post-synaptic location for F1-F3 or B1-B3 CMuGs (Figure 6C-D; quantified in Figure 6E). We conclude that co-activity of CMuGs can arise from widely distributed and overlapping MN post-synaptic sites.

**Figure 6.**
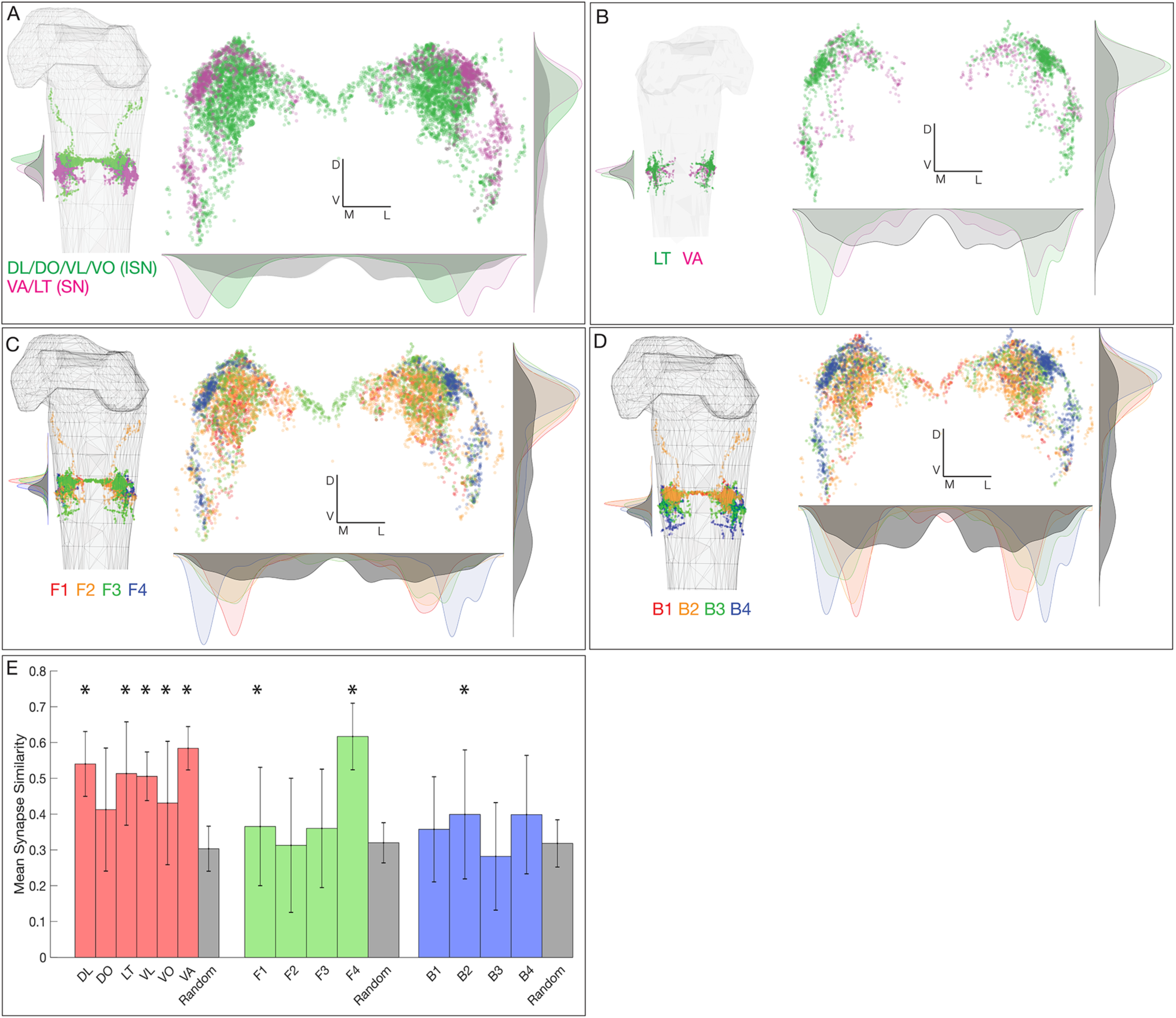
Motor neurons innervating SMuGs or CMuGs have post-synapses in distinct but overlapping regions of neuropil. (A-B) Spatial distributions of post-synaptic sites for MN pools innervating different SMuGs. Plots are 1D kernel density estimates for the A/P axis (A,B), or dorsoventral/mediolateral axes (A’/B’). Grey plots are density estimates for all post-synapses in A1 to illustrate the neuropil boundaries. (A) ISN (DL/DO/VL/VO) and SN (VA/LT) MN pools show segregation of inputs along mediolateral and anteroposterior axes. (B) VA and LT MN pools show segregation of inputs along mediolateral and anteroposterior axes. (C-D) Spatial distributions of post-synaptic sites for MN pools innervating different CMuGs. Plots are 1D kernel density estimates for the A/P axis (A,B), or dorsoventral/mediolateral axes (A’/B’). Grey plots are density estimates for all post-synapses in A1 to illustrate the neuropil boundaries. (C) Comparison of F4 MN pools with any of F1-F3 MN pools show segregation of inputs along mediolateral and anteroposterior axes. F1-F3 MN pools show a high degree of overlap. (D) Comparison of B4 MN pools with any of B1-B3 MN pools show segregation of inputs along mediolateral and anteroposterior axes. F1-F3 MN pools show a high degree of overlap. (E) Mean synapse similarity of spatial muscle groups and CMuGs. Each bar represents the mean pairwise synapse similarity of MNs in a given group, with self-similarity excluded. Random bars represent the mean synapse similarity of 100 groups of randomly selected neurons. Random group size was set as the mean size of spatial muscle groups, forward CMuGs, or backward CMuGs respectively. Asterisks denote p-values of less than .05 when compared to the random group.

### Premotor neuron pools can innervate individual CMuGs

In mammals there are dedicated PMN pools that innervate functionally related muscles (reviewed in Arber 2017; Arber and Costa 2018). Here we ask if the same “labeled line” connectivity is observed in the *Drosophila* motor connectome. There are six distinct SMuGs, and four CMuGs for forward and backward locomotion. We asked if there are groups of PMNs that are dedicated to individual SMuGs or CMuGs. We observed that many PMNs provided highly enriched synaptic input to MNs innervating a single SMuG (Figure 7A). For example, the PMNs shown in gray strongly prefer to form synapses with MNs innervating DL muscles, whereas PMNs shown in blue prefer to form synapses with MNs innervating LT muscles (Figure 7A). Despite this bias, no PMN exclusively formed synapses onto MNs in a single SMuG, nor did we observe any PMN that was strongly connected to MNs in all SMuGs (Figure 7A). We also observed a few PMNs that preferentially innervated a single forward CMuG (Figure 7B). For example, PMNs in magenta strongly preferred MNs innervating CMuG F2, PMNs in green were enriched for MNs targeting CMuG F3, and PMNs in dark blue show enriched connectivity to CMuG F1 and/or F2 (Figure 7B). This occurs despite the lack of MN post-synapse clustering described above for the F1-F3 CMuGs. Similarly, some PMNs showed enriched connectivity to a single backward CMuGs (Figure 7C). For example, PMNs in orange preferred MNs innervating CMuG B2, PMNs in light green strongly preferred MNs innervating CMuG B3, and PMNs in light blue show enriched connectivity to CMuG B3 and/or B4 (Figure 7C). Finally, we identified PMNs (e.g. A18b2, A27h) with strong connections to a single forward and/or backward CMuG, and sparser connections to other CMUGs. To investigate the functional significance of the observed PMN-MN CMuG connectivity, we performed dual calcium imaging on MNs innervating CMuGs F1/F2 and the A27h PMN with enriched connectivity to the CMuG F3. We found that as predicted by connectivity, CMuG F1/F2 MNs fired ahead of A27h, which is preferentially innervating MNs in CMuG F3 (Figure 7D). We conclude that there are PMNs preferentially innervating specific CMuGs, although there are many PMNs that innervate multiple CMuGs.

**Figure 7.**
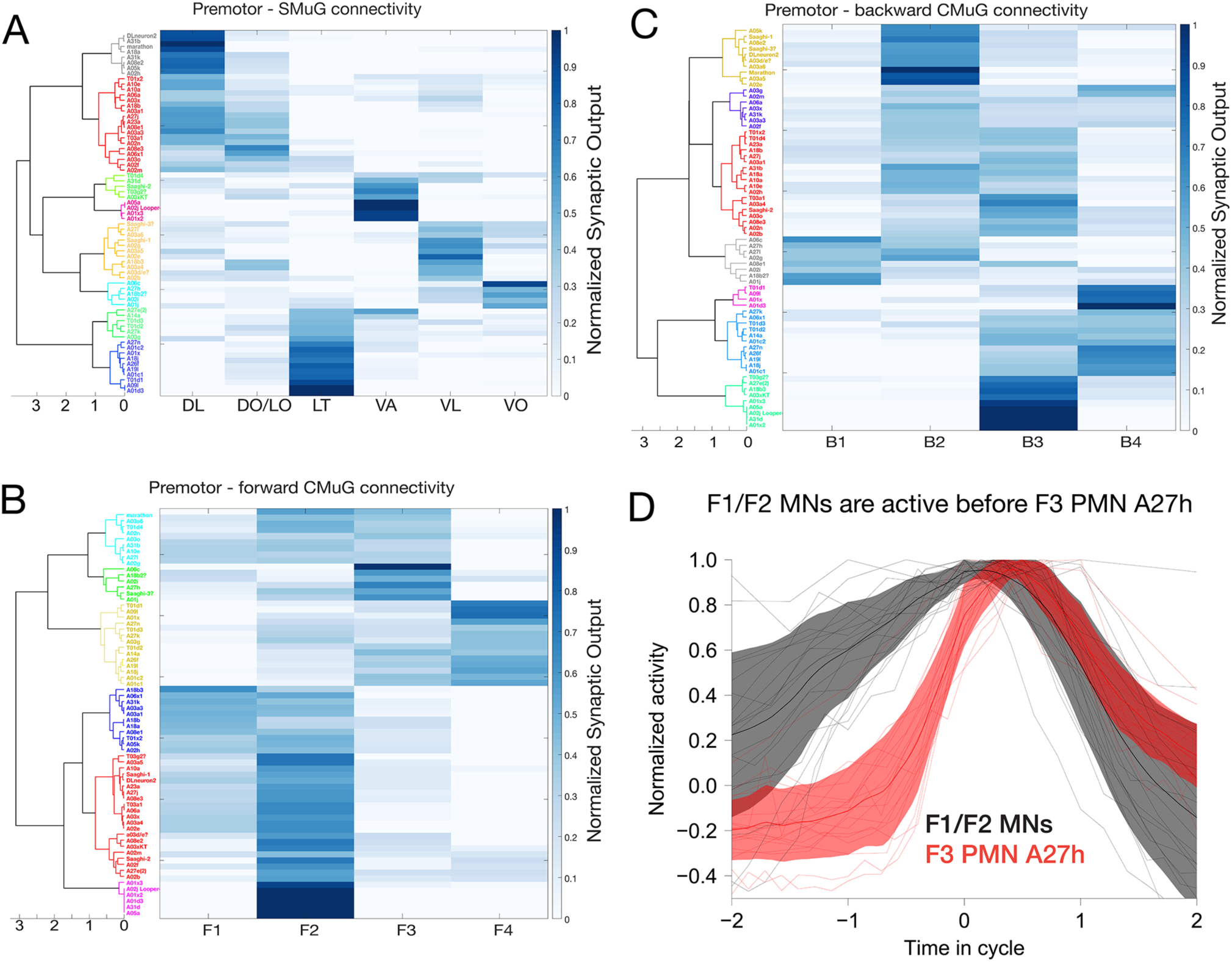
PMN pools preferentially connected to individual SMuGs and CMuGs. (A-C) Heirarchical clustering of PMNs based on their connectivity to MNs of the same spatial muscle group (A), forward CMuG (B), or backward CMuG (C). Heat maps represent the normalized weighted-synaptic output of a given left/right pair of PMNs onto grouped left/right pairs of MNs. PMN output strength was weighted to account for the number of inputs onto a given MN. For forward and backward CMuGs where not all MNs are represented, normalization was done after removing PMN inputs to the missing MNs. (A) Pools of PMNs show enriched connectivity to SMuGs (dark blue). (B) Pools of PMNs show enriched connectivity to F1-F4 CMuGs (dark blue). (C) Pools of PMNs show enriched connectivity to B1-B4 CMuGs (dark blue). (D) Dual color calcium imaging of jRCaMP1b in A27h (red) and GCaMP6m in U1-U5 MNs (black). Consistent with predictions from the connectome, U1-U5 MNs (CMuG F1/2) are activated before A27h (CMuG F3) during forward locomotion. Red and dark error bars (ribbons) represent the standard deviation of the average neuronal activity. Genotype: *CQ-lexA/+; lexAop-GCaMP6m/R36G02-Gal4 UAS-jRCaMP1b*.

### Neural circuit motifs predicted to generate distinct motor behaviors

In the previous section, we identified PMNs with enriched connectivity to specific forward and/or backward CMuGs. Here we further focus on PMN-MN connectivity and identify circuit motifs that are consistent with the observed CMuG timing, providing candidate motifs for functional studies. We highlight <u>intra</u>segmental motifs that could produce the observed phase delays between the four CMuGs in a single segment, <u>inter</u>segmental motifs that could produce the sequential activation of a specific CMuG in adjacent segments, motifs that could change the relative activation order of a MN/muscle between forward and backward locomotion, and a motif that could drive motor output initiating escape behavior.

#### Intrasegmental phase delays during forward or backward locomotion

Interactions between PMNs are likely to establish the phase delay between the four CMuGs as the motor wave moves across a segment. We used connectome and neurotransmitter data to identify PMN circuit motifs consistent with the observed CMuG intrasegmental phase relationships. First, we identified a disinhibition motif that could generate a phase delay between CMuGs F2 and F3/4. A02i and A14a (preferentially connected to CMuG F3/4) synapse onto A02e (preferentially connected to CMuG F1/F2). All of these PMNs are inhibitory, and thus this motif may disinhibit F1/F2 while inhibiting F3/4, producing a phase delay between F2 and F3/4 (Figure 8A). This confirms and extends previous work showing that A14a creates a phase delay between LO1 (CMuG F2) and LT1 (CMuG F4) (Zwart et al. 2016). We also observed a feedforward excitatory motif that could help synchronize MN activity within individual CMuGs (Figure 8B) and a feedforward inhibitory motif that could generate a phase delay between early CMuGs (F1/F2 or B1/B2) and late CMuGs (F4 or B4)(Figure 8B). Furthermore, we found another feedforward excitatory motif that could explain how MNs innervating different CMuGs show overlapping peak activity later in the contraction cycle within a segment. The previously described excitatory PMN A27h (Fushiki et al. 2016; Carreira-Rosario et al. 2018) connects to two excitatory PMNs, A18b2 and A18b3, which in turn connect to MNs innervating F1-F4 CMuGs. A27h has enriched connectivity to F3 CMuG which fires with a delay after F1/F2 MNs, and thus when A27h activates CMuG F3, it also activates A18b2 and A18b3, hence ensuring continued excitation to earlier CMuGs F1/F2 (Figure 8C). These motifs provide testable hypotheses for how specific phase relationships between CMuGs are generated by PMNs.

**Figure 8.**
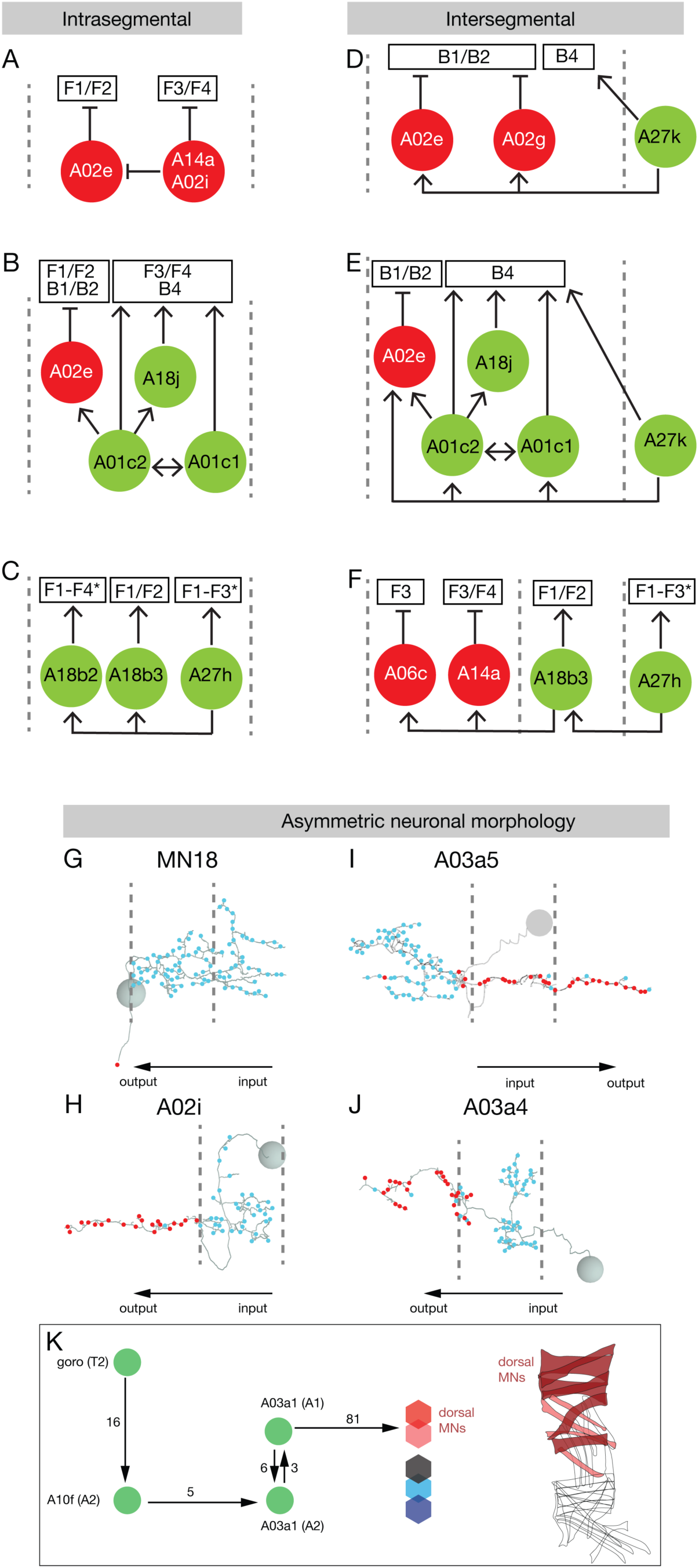
Neural circuit motifs consistent with MN phase delays and escape behavior. (A) Intrasegmental phase delay motif. Disinhibitory motif that could induce a phase delay between CMuG F1/2 and F3/4. Green circles and arrows indicate excitatory connection, red circles and T-bars indicate inhibitory connection. Anterior is to the left; segment boundaries are dashed lines determined by the position of aCC MN cell bodies. (B) Intrasegmental feedforward and recurrent excitatory motifs between green PMNs. This motif may ensure that F3/F4 and B4 CMuGs receive a sufficient period of excitation from PMNs. Intrasegmental feedforward inhibitory motif between A01c2 and A02e PMNs. This motif could exert inhibition to F1/F2 and B1/B2 CMuGs while simultaneously providing excitation to F3/F4 and B4 CMuGs. Labels as in A. (C) Intrasegmental feedforward excitatory motifs between PMNs. This motif may ensure that MNs innervating F1-F4 CMuGs receive simultaneous excitation late in the contraction cycle where all CMuGs show overlapping peak activity. (D) Intersegmental phase delay motif. Feedforward inhibitory motif that could induce a phase delay between CMuG F1/2 and F4 in adjacent segments. Labels as in A. (E) Intersegmental feedforward excitatory motif between A27k and A01c1/c2 PMNs with potential role during backward crawling. This motif may ensure that MNs innervating B4 CMuG receive temporally long enough excitation from PMNs. Intersegmental feedforward inhibitory motif between A27k and A02e PMNs with potential role in temporal segregation of B1/B2 from B4. (F) Intersegmental feedforward excitatory motif between A27h and A18b3 PMNs with potential role during forward crawling. Based on this motif, while A27h excites F1-F3 CMuGs in its own segment, it also activates F1/F2 CMuGs of the next anterior segment via A18b3 PMN. * A27h is heavily connected to F3 MNs and sparsely connected to F1/F2 MNs. Intersegmental feedforward inhibitory motif between A18b3 and A06c/A14a PMNs with potential role during forward crawling. Based on this motif, while A18b3 excites F1-F2 CMuGs in its own segment, it also inhibits F3/F4 CMuGs of the next anterior segment via A06c/A14a PMNs. This inhibition may relax the F3/F4 muscles ahead of upcoming forward motor wave. (G-J) Neuronal asymmetry along the anterior-posterior axis may contribute to intersegmental phase delays of individual CMuGs. (G) MN18 has asymmetric posterior dendrites that could be activated earlier during forward locomotion than during backward locomotion. (H) PMN A02i has an asymmetric anterior axon that could inhibit target MNs earlier during forward locomotion than during backward locomotion. (I) PMN A03a5 has an asymmetric posterior axon that could induce target MNs earlier during backward locomotion than during forward locomotion. (J) PMN A03a4 has an asymmetric anterior axon that could induce target MNs earlier during forward locomotion than during backward locomotion. Anterior, left; segment boundary, dashed line. (K) Multi-synaptic pathway from the escape-response inducing command neuron (goro) to motor neurons innervating dorsal muscle.

#### Intersegmental phase delays during forward or backward locomotion

We identified both feedforward excitation and feedforward inhibition motifs that could explain the sequential activation of a specific CMuG in adjacent segments during peristaltic motor waves. The excitatory PMN A27k (preferentially connected to CMuG B4) is involved in a feedforward inhibitory circuit in which it excites the inhibitory local PMNs A02e and A02g (preferentially connected to CMuG B1/B2). This motif could terminate B1/B2 activity and allow B3/4 activity as the contraction wave moves posteriorly (Figure 8D). A27k also synapses in the next anterior segment with the trio of excitatory neurons described above (A01c1, A01c2, and A18j) which are preferentially connected to CMuG B4, as well as with the inhibitory A02e PMN connected to CMuGs B1/B2. Thus, when an anterior-to-posterior backwards wave stimulates A27k, it results in A27k activating PMNs that trigger CMuG B4 as well as the PMN A02e that inhibits CMuGs B1/B2 in the next anterior segment (Figure 8E). Furthermore, we found feedforward excitatory and inhibitory motifs that could explain how different CMuGs in the adjacent segments are coordinated. A27h excites A18b3 in the next anterior segment to move the contraction wave forward, while A18b3 excites the inhibitory neurons A06c/A14a to prevent premature activation of neurons in CMuG F3/4 in the next adjacent segment (Figure 8F).

#### Distinct MN phase relationships during forward and backward locomotion

As described earlier, forward and backward locomotion do not simply correspond to the same CMuG1-4 timing in reverse order (Figure 5). For example, MN18 is active in the early during forward locomotion, but late during backward locomotion (Figure 5). Interestingly, MN18 dendrites extend uni-directionally to the adjacent posterior segment (Figure 8G) (Landgraf et al. 1997). The asymmetric morphology of MN18 is likely to explain its earlier activation in forward locomotion, as the peristaltic wave would engage its dendrites earlier. Similar anterior/posterior asymmetry was observed in nine PMNs: A02i, A02j, A03a4, A18b2, A18b3, A26f, A27k that all had axons extending 1-2 segments anterior of the cell body and dendrites, and A01j and A03a5 that had axons projecting 1-2 segments posterior to the cell body and dendrites (Figure 8I-J; Figure 3 – supplement 1). The anterior/posterior asymmetry in axon projection of these PMNs suggests that they may exert temporally distinct effects on their target MNs during forward versus backward locomotion, raising the possibility that they may have different roles in these two behaviors. For example, when the cholinergic A03a5 PMN (Takagi et al. 2017) becomes active during a forward or backward wave, its posteriorly-projecting (descending) axons will excite MNs located in its own segment as well as further posterior segments. As a result, given the opposite direction of wave propagation in backward and forward locomotion, MN targets of A03a5 will receive excitatory inputs earlier during backward than forward locomotion (Figure 8I). We conclude that PMN/MN morphological asymmetry may contribute to the differential timing of muscle activation during between forward and backward locomotion.

#### A motor pathway for escape behavior?

*Drosophila* larvae escape from nociceptive stimuli by performing a three step escape behavior: c-bend, lateral rolling, and fast forward locomotion. This behavior can be triggered by optogenetic activation of the descending neuron “gorogoro,” and the pathway from nociceptors to gorogoro has been well-characterized (Ohyama et al. 2015; Takagi et al. 2017; Burgos et al. 2018). However, nothing is known about how gorogoro triggers the escape motor program. Here we find a candidate motor circuit that may initiate C-bending. We find that gorogoro connects via A10f to the PMN A03a1, which specifically innervates dorsal body wall muscles (Figure 8K). The polarity of A10f and A03a1 are unknown, but if they are both excitatory, then gorogoro could specifically activate dorsal muscles, which we have previously shown is sufficient to induce larval bending (Clark et al. 2016).

### Proprioceptor-premotor connectivity predicts a role in temporal focusing of motor neuron activity

Proprioception is the sensory modality with the most direct pathway to motor neurons (Imai and Yoshida 2018). Our PMN/MN connectome provides an opportunity to determine whether proprioceptors have direct or indirect connectivity to PMN/MNs, particularly since all six body wall proprioceptors have been previously traced in the TEM volume (Heckscher et al. 2015; Ohyama et al. 2015). Recent work has shown that five proprioceptors (ddaE, ddaD, vpda, dmd1, and vbd) are activated by muscle contraction and fire sequentially during forward locomotion; the sixth proprioceptor, dbd, is active during muscle relaxation (He et al. 2019; Vaadia et al. 2019). Genetic inhibition of proprioceptor function results in slower crawling due to prolonged muscle contraction during each peristaltic wave, which has led to the model that proprioceptors send a “mission accomplished” signal to terminate muscle contraction (Hughes and Thomas 2007).

We examined the relationship between proprioceptors and PMNs to identify circuit motifs that could generate a “mission accomplished” signal (from ddaE, ddaD, vpda, dmd1, and vbd) or terminating the signal (from dbd). We found strong connectivity between proprioceptor neurons and PMNs, but surprisingly little direct connectivity to MNs (Figure 9A). Note that less than 20% of the proprioceptor pre-synaptic sites are targeted to the PMNs we have characterized (Figure 9B), indicating that there may be additional PMNs yet to be characterized, and/or that proprioceptors preferentially connect to pre-PMNs. We find that **vbd** gives excitatory input to the inhibitory PMNs A02e/A02g (Figure 9C) or A02k (Figure 9D), which connect to MNs active throughout the contraction cycle. Thus, this circuit motif would contribute to a “mission accomplished” signal terminating muscle contraction and speeding locomotion. Similarly, **dmd1**/**ddaD** activation of the inhibitory PMN A27j could also send a “mission accomplished” signal (Figure 9E). In support of the functional significance of this motif, we find that A27j is rhythmically active during both forward and backward fictive locomotion in isolated brains, although we were unable to determine its phase-relationship with proprioceptors in this reduced preparation (data not shown). A similar motif contains **ddaE** and other proprioceptors, which innervate the inhibitory PMN A02b, connecting to the excitatory premotor neurons A03a6 and A18j (Figure 9F). In contrast, the other ventral proprioceptor, **vpda**, directly activates the excitatory premotor neuron A27h, which may be required to maintain MN activity long enough to drive complete muscle contraction during a crawl cycle (Figure 9G). Finally, the stretch-activated **dbd** proprioceptor is found in a motif which could activate the next muscle contraction cycle (Figure 9H). We conclude that all six proprioceptor neurons have direct or indirect connection to PMN circuit motifs that could allow them to promote efficient propagation of the muscle contraction wave during forward locomotion.

**Figure 9.**
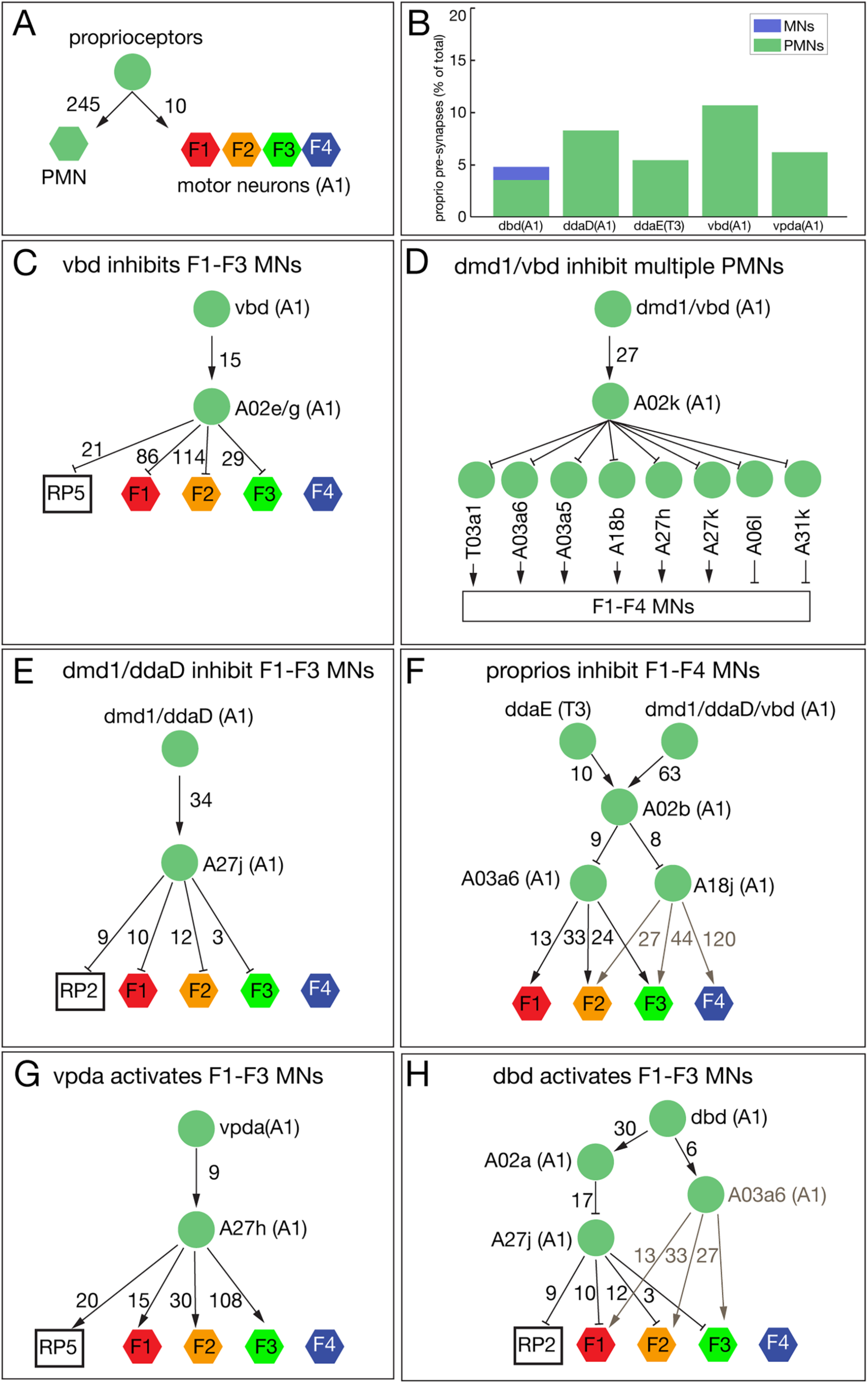
Proprioceptor-premotor motifs involved in locomotor wave propagation. (A-B) Synapse numbers between the six excitatory proprioceptors (dbd, vbd, vpda, dmd1, ddaD, and ddaE), PMNs, and MNs. PMN intra-group connectivity not shown. (C-H) Motifs containing the proprioceptors active during segment contraction (vbd, dmd1, ddaD, ddaE, vpda) and segment stretch (dbd). Arrowheads and T bars indicate excitatory and inhibitory connections, respectively. Number of synapses are shown next to each connection. F1-F4 indicates MNs innervating F1-F4 CMuGs. Type-Is MNs RP5 and RP2 are shown separately due to their broad connectivity. (C) Motif where vbd inhibits MNs in CMuGs F1-F3. (D) Motif where vpda activates MNs in CMuGs F1-F3. (E) Motif where dmd1/ddaD inhibit MNs in CMuGs F1-F3. (F) Motif where dmd1/ddaD/ddaE and vbd indirectly inhibit MNs in CMuGs F1-F4. (G) Motif where dmd1/vbd indirectly inhibit MNs in CMuGs F1-F4, and disinhibit A06l/A31k MNs. (H) Motif where the stretch proprioceptor, dbd, disinhibits MNs via A02a/A27j, and also directly activates MNs via the excitatory PMN A03a6.

### Modeling interactions among PMNs that generate sequential MN activation

Recurrent interactions among PMNs have been shown to control the timing of the muscle outputs of central pattern generator circuits in a variety of organisms (Marder and Bucher 2001; Grillner 2003). We hypothesized that these types of interactions are responsible for the timing of muscle activation during *Drosophila* larval forward and backward crawling. To assess whether the reconstructed PMN connectome is capable of producing the observed timing of MN/muscle activation, we developed a recurrent network model of two adjacent segments. Previous models have focused on wave propagation during forward and backward crawling by modeling the average activity of excitatory and inhibitory subpopulations in each segment (Gjorgjieva et al. 2013; Pehlevan et al. 2016). Access to the detailed connectivity of selected pre-PMNs, PMNs and MNs, as well as knowledge of the activation patterns of different CMuGs, allowed us to develop a substantially more detailed model whose circuitry was constrained to match the TEM reconstruction. For PMNs whose neurotransmitter identity we could determine, we also constrained the signs (excitatory or inhibitory) of connection strengths in the model. The firing rates of PMNs and MNs were modeled as simple threshold-linear functions of their synaptic inputs, and model parameters were adjusted to produce target MN patterns of activity that matched the sequences identified during forward and backward crawling. These patterns were assumed to be evoked by external command signals, representing descending input to the PMNs, that differed for forward and backward crawling but did not themselves contain information about the timing of individual muscle groups. We also constrained the activity of two PMNs, A18b and A27h, that are known to be specifically active during backward and forward locomotion, respectively (Fushiki et al. 2016; Carreira-Rosario et al. 2018). We found that, although the connectivity among PMNs within a segment is sparse (roughly 7% of all possible pairwise connections), the observed connections are nonetheless sufficient to generate appropriately timed MN activity for the two distinct behaviors (Figure 10A,B; Figure 10 – Figure supplement 1; see Methods). As has been described previously in other pattern-generating systems (Prinz et al. 2004), there is a space of models that is capable of producing the observed activity. We therefore analyzed the activity of neurons in an ensemble of models. In the models, distinct sequences of PMN activity for forward and backward locomotion tile the period of time over which MNs are active (Figure 10; Figure 10C – Figure supplement 1). These sequences give rise to the distinct timing of MN activation during each behavior. With the exception of *C. elegans* models (Karbowski et al. 2008; Macosko et al. 2009; Wen et al. 2012a; Izquierdo and Beer 2013; Izquierdo et al. 2015; Kunert et al. 2017; Rakowski and Karbowski 2017), the networks constructed here represent the first models of the neural circuitry underlying a timed motor behavior whose connectivity has been constrained by a synaptic wiring diagram.

**Figure 10.**
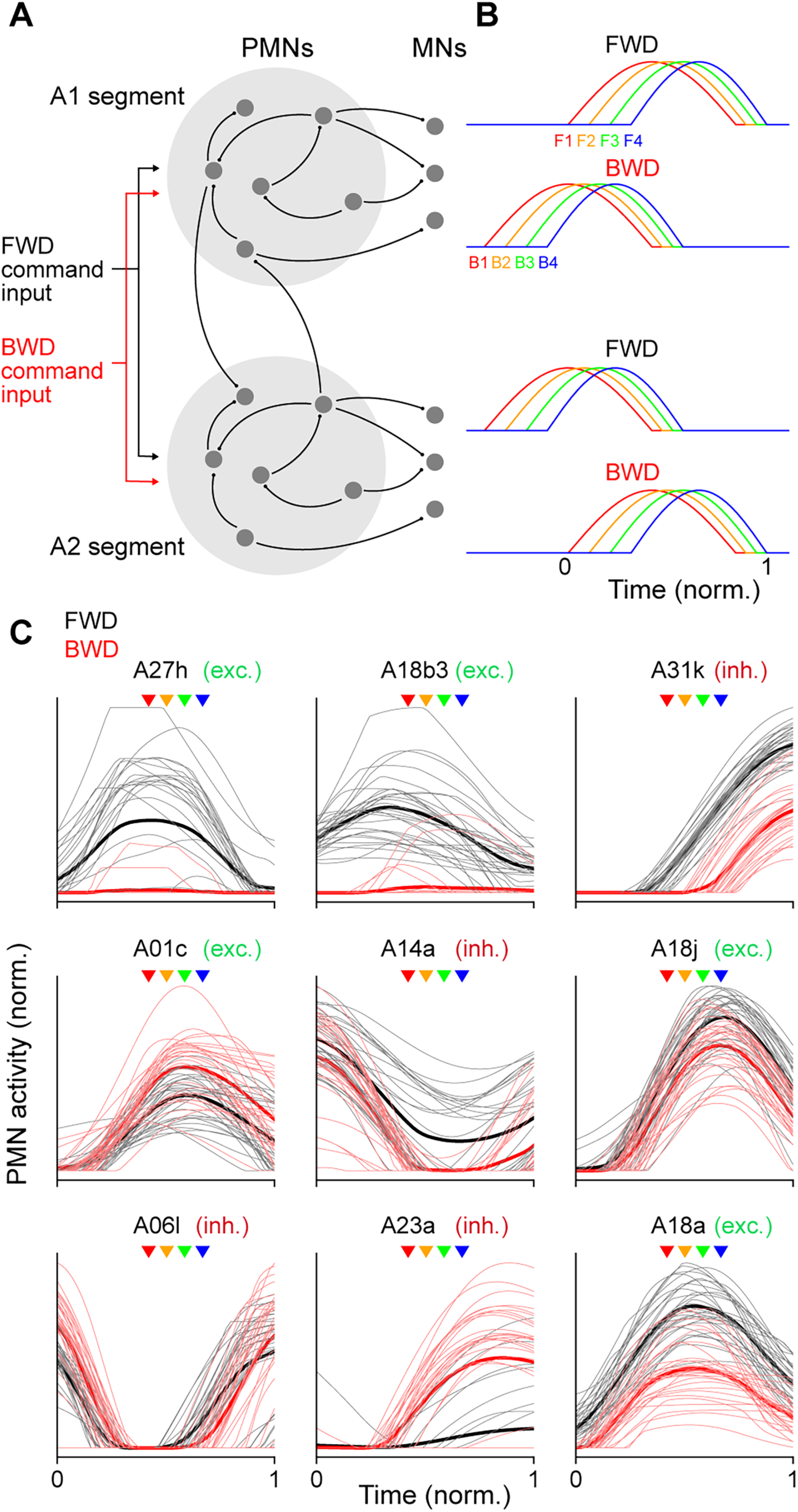
Recurrent network model generating sequential MN activity. (A) The PMN and MN network of the A1 and A2 segments was modeled using connectivity taken from the EM reconstruction. Connections within each segment (light gray circles) are identical. The network was optimized using gradient descent to produce a sequential pattern of activity in the MNs (MNs) when a tonic external command input for forward (forward, black) or backward (backward, red) locomotion was applied. (B) The network in A was optimized to produce an appropriate sequential activity pattern of CMuGs during forward and backward crawling. The direction of propagation from the posterior (A2) to anterior (A1) segment or vice versa differs for forward and backward crawling. To compare PMN activity relative to MN activation, time is measured in units normalized to the onset and offset of MN activity in a segment (bottom right). (C) Normalized activity of a subset of PMNs in the model during forward and backward crawling. Thick lines denote averages over the ensemble of models generated. Time is measured relative to MN onset and offset as in B. Arrowheads denote the peak activation onset time for the MNs innervating different CMuGs (color key as in panel B); exc, excitatory; inh, inhibitory.

### Experimental validation of the recurrent network model

Next we asked if the sequences of PMN activity predicted by the model are consistent with prior experimentally determined activity patterns. In our model, A14a shows high activity during CMuG F1 and is inactive during CMuG F4 (Figure 10C). Experimental data show that A14a is inhibitory and is active during CMuG F1; and blocking A14a activity removes the contraction delay between muscles in CMuG F1 and F4 (Zwart et al. 2016), validating our model. In our model, A18b3 and A18a are both active during forward locomotion, but only A18a is active during backward locomotion (Figure 10C). Experimental data show that A18a and A18b3 are active precisely as proposed in our model (Hasegawa et al. 2016). Furthermore, our model predicts the cholinergic A18j and A01c PMNs are active late in the crawl cycle during CMuG F4, which is supported by experimental data on these neurons (where they were called eIN1,2; Zwart et al. 2016).

To provide new, additional experimental tests of our model, we performed dual color calcium imaging on previously uncharacterized PMNs A31k, A06l, and A23a. First, our model predicted that both A31k and A06l neurons would be active with a phase delay relative to MN activity (Figure 10C; Figure 10 – supplement 1). To determine the phase-relationship between A31k and MNs, we expressed GCaMP6m in a subset of MNs and jRCaMP1b in A31k. Dual color calcium imaging data revealed that the A31k activity peak coincides with a decline of activity in MNs innervating early CMuGs (Figure 11A,B), validating our model. Second, we focused on the GABAergic inhibitory A31k and A06l neurons. Our model predicts that both PMNs show concurrent, rhythmic activity during forward and backward locomotion (Figure 10 – supplement 1). We expressed GCaMP6m in both neurons, which we could distinguish based on different axon projections, and found that they showed concurrent, rhythmic activity (Figure 11C,D), and thus both neurons show a delayed activation relative to MNs. Our third experimental test focused on the GABAergic A23a PMN (Schneider-Mizell et al. 2016). Our model predicted that A23a was active earlier during backward locomotion than forward locomotion (Figure 10C). We expressed GCaMP6m in a subset of MNs and jRCaMP1b in A23a, and validated the prediction of our model (Figure 11E,F). We conclude that our model accurately predicts many, but not all (see Discussion), of the experimentally determined PMN-MN phase relationships.

**Figure 11.**
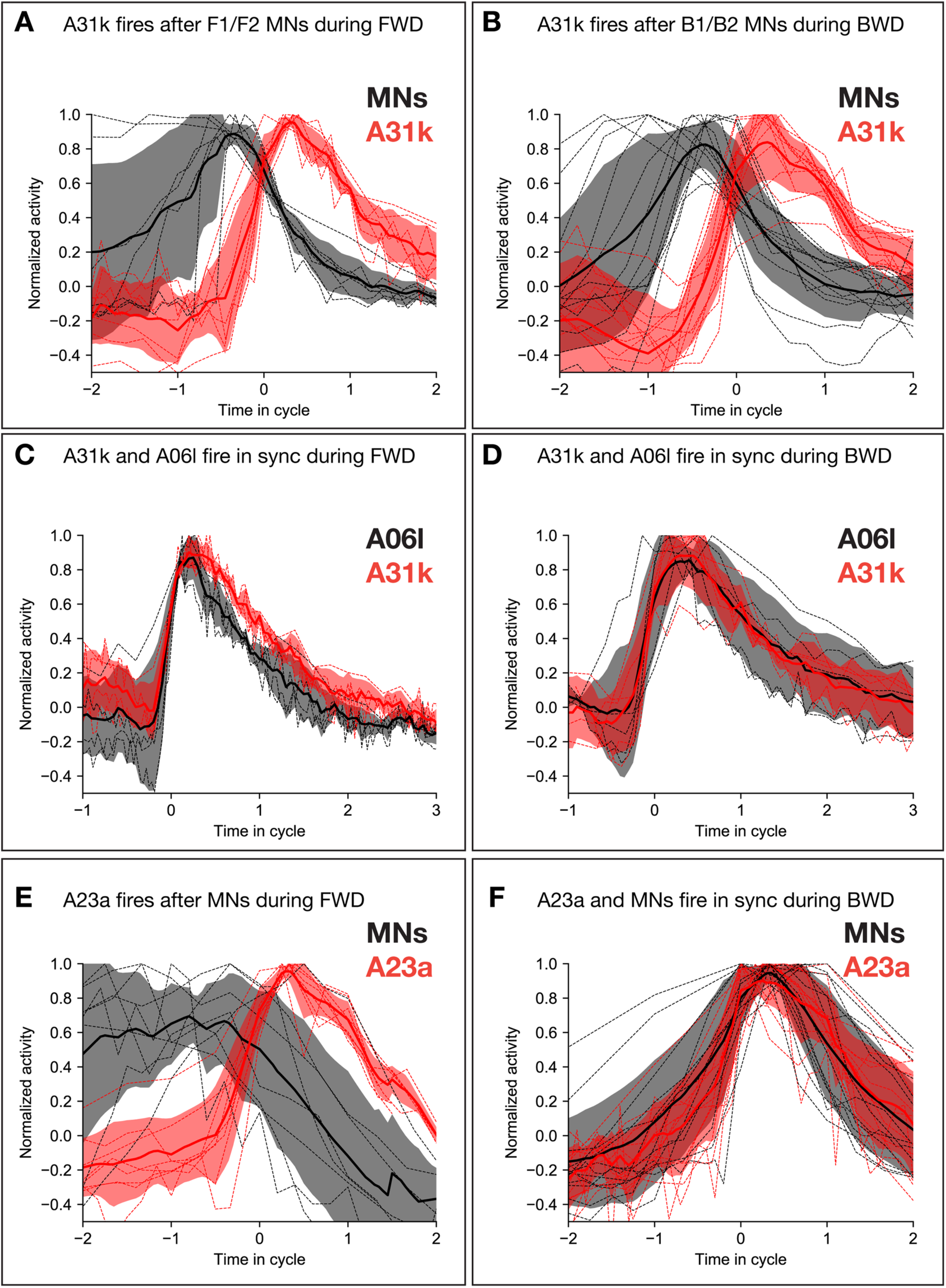
Calcium imaging of A31k/A06l/A23a PMNs and their target MNs validates the activity pattern predicted by recurrent modeling. (A-B) Dual color calcium imaging of jRCaMP1b in A31k (red) and GCaMP6m in MNs (black). Consistent with the recurrent model predictions, A31k fires with a delay after its post-synaptic MNs in both forward (A) and backward (B) waves. Red and dark error bars (ribbons) represent the standard deviation of the average neuronal activity. Genotype: *CQ-lexA/+; lexAop-GCaMP6m/R87H09-Gal4 UAS-jRCaMP1b*. (C-D) Single color calcium imaging of jRCaMP1b in A31k (red) and A06l (black). Consistent with the recurrent model predictions, A31k and A06l show synchronous activity patterns during forward (C) and backward waves (D). Red and dark error bars (ribbons) represent the standard deviation of the average neuronal activity. Genotype: *R87H09-Gal4 UAS-jRCaMP1b* (E-F) Dual color calcium imaging of jRCaMP1b in A23a (red) and GCaMP6m in MNs (black). Consistent with the recurrent model predictions, A23a fires with a delay after its post-synaptic MNs in forward (E) and synchronously with them in backward (F) waves. Red and dark error bars (ribbons) represent the standard deviation of the average neuronal activity. Genotype: *CQ-lexA/+; lexAop-GCaMP6m/R78F07-Gal4 UAS-jRCaMP1b*.

## Discussion

It is a major goal of neuroscience to comprehensively reconstruct neuronal circuits that generate specific behaviors, but to date this has been done only in *C. elegans* (Karbowski et al. 2008; Macosko et al. 2009; Izquierdo and Beer 2013; Izquierdo et al. 2015; Kunert et al. 2017; Rakowski and Karbowski 2017). Recent studies in mice and zebrafish have shed light on the overall distribution of PMNs and their connections to several well-defined MN pools (Eklof-Ljunggren et al. 2012; Kimura et al. 2013; Bagnall and McLean 2014; Ljunggren et al. 2014). However, it remains unknown if there are additional PMNs that have yet to be characterized, nor do we have any insight into potential connections between PMNs themselves, which would be important for understanding the network properties that produce coordinated motor output. In the locomotor central pattern generator circuitry of leech, lamprey, and crayfish, the synaptic connectivity between PMNs or between PMNs and other interneurons are known to play critical roles in regulating the swimming behavior (Brodfuehrer and Thorogood 2001; Grillner 2003; Kristan et al. 2005; Mullins et al. 2011; Mulloney and Smarandache-Wellmann 2012; Mulloney et al. 2014). However, it is difficult to be certain that all the neural components and connections of these circuits have been identified. Thus, the comprehensive anatomical circuitry reconstructed in our study provides an anatomical constraint on the functional connectivity used to drive larval locomotion; all synaptically-connected neurons may not be relevant, but at least no highly connected local PMN is absent from our analysis.

Our results confirm and significantly extend previous studies of *Drosophila* larval locomotion. For example, a recent study (Zwart et al. 2016) has shown that the GABAergic A14a inhibitory PMN (also called iIN1) selectively inhibits MNs innervating muscle 22/LT2 (CMuG F4), thereby delaying muscle contraction relative to muscle 5/LO1 (CMuG F2). We extend this study by showing that A14a also disinhibits MNs in early CMuGs F1/2 via the inhibitory PMN A02e. Thus, A14a both inhibits late CMuGs and disinhibits early CMuGs. In addition, previous work has suggested that all MNs receive simultaneous excitatory inputs from different cholinergic PMNs (Zwart et al. 2016). However, our dual calcium imaging data of the A27h excitatory PMN shows that it is active during CMuG F4 and not earlier. Therefore, MNs may receive temporally distinct excitatory inputs, in addition to the previously reported temporally distinct inhibitory inputs. We have identified dozens of new PMNs that are candidates for regulating motor rhythms; functional analysis of all of these PMNs is beyond the scope of this paper, particularly due to the additional work required to screen and identify Gal4/LexA lines selectively targeting these PMNs, but our predictions are clear and testable when reagents become available.

We show that MNs innervating a single SMuG belong to more than one CMuG, therefore SMuGs do not generally match CMuGs. This could be due to the several reasons: (i) MNs in each SMuGs receive inputs from overlapping but not identical array of PMNs (Table 2). (ii) Different MNs in the same SMuG receive a different number of synapses from the same PMN (Supplementary Table 1). (iii) MNs in the same SMuG vary in overall dendritic size and total number of post-synapses (Supplementary Table 1), thereby resulting in MNs of the same SMuGs fall into different CMuGs.

We demonstrate that during both forward and backward crawling, most of longitudinal and transverse muscles of a given segment contract as early and late groups, respectively. In contrast, muscles with oblique or acute orientation often show different phase relationships during forward and backward crawling. Future studies will be needed to provide a biomechanical explanation for why oblique muscles – but not longitudinal or transverse muscles – need to be recruited differentially during forward or backward crawling. Also, it will be interesting to determine which spatial muscle groups (e.g. either or both VOs and VLs) are responsible for elevating cuticular denticles during propagation of the peristaltic wave in forward and backward crawling; if the VOs, it would mean that lifting the denticles occurs at different phases of the crawl cycle in forward and backward locomotion. Finally, understanding how the premotor-motor circuits described here are used to generate diverse larval motor behaviors will shed light on mechanisms underlying the multi-functionality of neuronal circuits.

A recent study has reported that proprioceptive sensory neurons (dbd, vbd, vpda, dmd1, ddaE, and ddaD) show sequential activity during forward crawling. dbd responds to stretching and whereas the other five classes are activated by muscle contraction (Vaadia et al. 2019). All proprioceptors show connectivity to the tier of PMNs we describe here, and we have identified circuit motifs that are consistent with the observed timing and excitatory function of each proprioceptor neuron (Figure 8). It will be of great interest perform functional experiments to test these anatomical circuit motifs for functional relevance.

Our recurrent network model accurately predicts the order of activation of specific PMNs, yet many of its parameters remain unconstrained, and some PMNs may have biological activity inconsistent with activity predicted by our model. Sources of uncertainty in the model include incomplete reconstruction of inter-segmental connectivity and descending command inputs, the potential role of gap junctions (which are not resolved in the TEM reconstruction), as well as incomplete characterization of PMN and MN biophysical properties. Recent studies have suggested that models constrained by TEM reconstructions of neuronal connectivity are capable of predicting features of neuronal activity and function in the *Drosophila* olfactory (Eichler et al. 2017) and visual (Takemura et al. 2013; Tschopp et al. 2018) systems, despite the unavoidable uncertainty in some model parameters (Bargmann and Marder 2013). Similarly, for the locomotor circuit described here, we anticipate that the addition of model constraints from future experiments will lead to progressively more accurate models of PMN and MN dynamics. Despite it’s limitations, the ability for the PMN network to generate appropriate muscle timing for two distinct behaviors in the absence of any third-layer or command-like interneurons suggests that a single layer of recurrent circuitry is sufficient to generate multiple behavioral outputs, and provides insight into the network architecture of multifunctional pattern generating circuits.

Previous work in other animal models have identified multifunctional muscles involved in more than one motor behavior: swimming and crawling in *C. elegans* (Pierce-Shimomura et al. 2008; Vidal-Gadea et al. 2011; Butler et al. 2015) and leech (Briggman and Kristan 2006); walking and flight in locust (Ramirez and Pearson 1988); respiratory and non-respiratory functions of mammalian diaphragm muscle (Lieske et al. 2000; Fogarty et al. 2018) unifunctional muscles which are only active in one specific behavior in the lobster *Homarus americanus* (Mulloney et al. 2014); swimming in the marine mollusk *Tritonia diomedea* (Popescu and Frost 2002); and muscles in different regions of crab and lobster stomach (Bucher et al. 2006; Briggman and Kristan 2008). Our single-muscle calcium imaging data indicates that all imaged larval body wall muscles are bifunctional and are activated during both forward and backward locomotion. It will be interesting to determine if all imaged muscles are also involved in other larval behaviors, such as escape rolling, self-righting, turning, or digging. It is likely that there are different CMuGs for each behavior, as we have seen for forward and backward locomotion, raising the question of how different CMuGs are generated for each distinct behavior.

## Methods

### Electron microscopy and CATMAID reconstructions

Neurons were reconstructed in CATMAID using a Google Chrome browser as previously described (Ohyama et al. 2015). Candidate PMNs were discarded if their maximum MN connectivity was ≤5 synapses (summed across the left and right hemispheres), where the neuron could not be traced due to gaps in the TEM volume, and a few neurons with massive arbors which were beyond our ability to trace. Figures were generated using CATMAID graph or 3D widgets combined with Adobe Illustrator (Adobe, San Jose, CA).

### Synapse spatial distributions and clustering

Synapse spatial distributions were generated using custom MATLAB scripts. Spatial distributions were determined using kernel density estimates with a 1 µm bandwidth. For cross-sectional spatial distributions, points were rotated −12 degrees around the Z-axis (A/P axis) in order to account for the slight offset of the EM-volume. For pre-synaptic sites (Figure 3D) polyadic synapses were weighted by their number of post-synaptic targets. Synapse similarity was calculated as described previously (Schlegel et al. 2016):

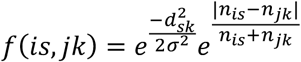

where *f(is,jk)* is the mean synapse similarity between all synapses of neuron *i* and neuron *j. d*_*sk*_ is the Euclidean distance between synapses *s* and *k* such that synapse *k* is the closest synapse of neuron *j* to synapse s of neuron *i*. σ is a bandwidth term that determines what is considered close. *n*_*is*_ and *n*_*jk*_ are the fraction of synapses for neuron *i* and neuron *j* that are within ω of synapse *s* and synapse *k* respectively. For MN inputs, σ = ω = 2 µm. Clustering was performed by using the average synapse similarity scores for the left and right hemisegments as a distance metric, and linkage was calculated using the average synapse similarity.

### Clustering analysis of PMN-MN connectivity

Hierarchical clustering was performed on the connectivity of PMNs to groups of MNs using weights inferred from the TEM. First, we summed left and right pairs of PMNs and MNs and divided the PMN-MN weights by the total number of inputs for a given MN, and summed the weighted connections for all MNs in a given muscle group. Then we normalized the vector of PMN-MN group connections before clustering. We used a Euclidean distance metric with wards linkage.

### Muscle GCaMP6f imaging, length measurement, and quantification

2% melted agarose was used to make pads with similar size: 25mm (W) X 50mm (L) X 2mm (H). Using tungsten wire, a shallow ditch was made on agarose pads to accommodate the larva. To do muscle ratiometric calcium imaging in intact animals, a first or second instar larvae expressing GCaMP6f and mCherry in body wall muscles were washed with distilled water, then moved into a 2% agarose pad on the slide. A 22 mm × 40 mm cover glass was put on the larva and pressed gently to gently constrain larval locomotion. The larva was mounted dorsolaterally or ventrolaterally to image a different set of muscles (dorsolateral mount excludes the most ventral muscles (15,16,17) whereas the ventrolateral mount excludes the dorsal-most muscles (1,2,9,10); imaging was done with a 10x objective on an upright Zeiss LSM800 microscope. We recorded a total of 38 waves (24 forward and 14 backward) from four different animals, and examined muscle calcium activity in two subsequent hemi-segments for each wave. Muscle length measurement was done using custom MATLAB scripts where muscle length was measured on a frame by frame basis. Calcium imaging data was also analyzed using custom MATLAB scripts. Due to movement artifacts, ROIs were updated on a frame by frame basis to track the muscle movement. ROIs that crossed other muscles during contraction were discarded. In no single preparation was it possible to obtain calcium traces for all 30 muscles. Instead, we used only preparations in which at least 40% of the muscles could be recorded. In order to align crawl cycles that were of variable time and muscle composition, we first produced a 2 dimensional representation of each crawl cycle using PCA. Crawl cycles were represented as circular trajectories away from, and back towards the origin (Figure 5 – figure supplement 1E,F) similar to what has been shown previously (Lemon et al. 2015). The amplitude, or linear distance from the origin, to a point on this trajectory correlated well with both the coherence of the calcium signals as well as the amplitude of the population. Thus, peaks in this 2D amplitude correspond with the time in which most muscles are maximally active, which we defined as the midpoint of a crawl cycle. It should be noted that the muscles used to generate two dimensional representations of crawl cycles were different for each crawl. While this means that each PCA trajectory is slightly different for each crawl cycle, we reasoned that because each experiment contained muscles from every CMuG, the peak amplitude in PCA space should still correspond to a good approximation of the midpoint of the crawl cycle. We defined the width of a crawl cycle as the width of this 2D peak at half-height (Figure 5 – figure supplement 1G). We aligned all crawl cycles to the crawl onset and offset (which we call 25% and 75% of the crawl cycle respectively) as defined by this width at half-height (Figure 5 – figure supplement 1H,I).

### Calcium imaging in neurons

For dual-color and single-color calcium imaging in fictive preps, freshly dissected brains were mounted on 12mm round Poly-D-Lysine Coverslips (Corning® BioCoat™) in HL3.1 saline (de Castro et al. 2014), which were then were placed on 25 mm × 75 mm glass slides to be imaged with a 40× objective on an upright Zeiss LSM-800 confocal microscopy. To simultaneously image two different neurons expressing GCaMP6m we imaged neuron-specific regions of interest (ROI). In addition, we imaged two neurons differentially expressing GCaMP6m and jRCaMP1b. Image data were imported into Fiji (https://imagej.net/fiji) and GCaMP6m and jRCaMP1b channels were separated. The ΔF/F_0_ of each ROI was calculated as (F-F_0_)/F_0_, where F_0_ was averaged over ∼1s immediately before the start of the forward or backward waves in each ROI.

### Antibody staining and imaging

Standard confocal microscopy, immunocytochemistry and MCFO methods were performed as previously described for larvae (Carreira-Rosario et al. 2018). Primary antibodies used: GFP or Venus (rabbit, 1:500, ThermoFisher, Waltham, MA; chicken 1:1000, Abcam13970, Eugene, OR), GFP or Citrine (Camelid sdAB direct labeled with AbberiorStar635P, 1:1000, NanoTab Biotech., Gottingen, Germany), GABA (rabbit, 1:1000, Sigma, St. Louis, MO), mCherry (rabbit, 1:1000, Novus, Littleton, CO), HA (mouse, 1:200, Cell Signaling, Danvers, MA), or V5 (rabbit, 1:400, Rockland, Atlanta, GA), Flag (rabbit, 1:200, Rockland, Atlanta, GA). Secondary antibodies were from Jackson Immunoresearch (West Grove, PA) and used according to manufacturer’s instructions. Confocal image stacks were acquired on Zeiss 710 or 800 microscopes. Images were processed in Fiji (https://imagej.net/Fiji), Photoshop, and Illustrator (Adobe, San Jose, CA). Brightness and contrast adjustments were applied to the entire image uniformly; mosaic images were assembled in Photoshop (Adobe, San Jose, CA).

### Theoretical modeling

We constructed a recurrent network representing the activity of PMNs, which we denote by the vector **p**, and of MNs, which we denote by the vector **m**. These vectors contain entries corresponding to neurons of two adjacent segments, so for example **p** = (**p**^1^, **p**^2^), where **p**^1^ and **p**^2^ represent activities of PMNs that innervate MNs of segments A1 or A2, respectively. The firing rate of PMN or MN *i* is a rectified-linear function of its input: 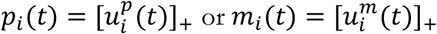, where [·]_+_ denotes rectification. The PMN input **u**^*p*^ follows the differential equation:

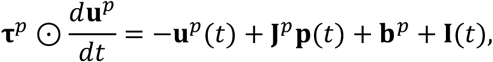

where 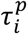 is the time constant of PMN 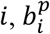 its baseline excitability, *I*_*i*_(*t*) its descending input from other regions, ⊙ denotes element-wise multiplication, and **J**^*p*^ is the connectivity matrix among PMNs. The descending input to the PMNs **I**(*t*) is represented as a pulse of activity: **I**(*t*) = **I**^*FWD*^ during FWD crawling, **I**(*t*) = **I**^*BWD*^ during BWD crawling, and **I**(*t*) = 0 otherwise.

MNs follow similar dynamics:

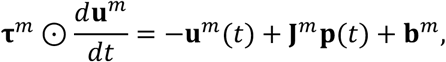

where **J**^*m*^ is the connectivity matrix from PMNs to MNs.

These matrices have block structures,

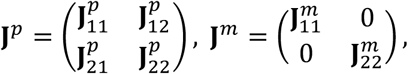

where 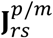 represents connections from segment *r* to segment *s*. We constrain 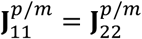. Due to lack of data, we have ignored PMNs that innervate MNs in multiple segments. We similarly constrain the entries of **τ**^*p*/*m*^ and **b**^*p*/*m*^ that correspond to the same neuron in different segments to be equal.

The model parameters (**J, b, τ, I**) are adjusted using gradient descent so that the MN activity **m** reproduces target patterns of activity during FWD and BWD crawling. These targets are defined for 6 s trials that contain one sequence of CMUG activation in each of the two segments. Time is discretized into 50 ms bins. At the beginning of each trial, **u**^*p*^ is initialized with random values from a truncated Gaussian distribution with standard deviation 0.1, and **u**^*m*^ is initialized to 0. A trial consists of sequential activity in each segment with a 1 s inter-segmental delay (Figure 9). Trials begin and end with 1 and 1.5 s of quiescence, respectively. Each MN’s target activity is given by a rectified cosine pulse of activity whose start and end times depend on the CMUG to which it belongs. The first CMUG is active for 2 s, and subsequent CMUGs activate with a delay of 0.25 s between each group and end with a delay of 0.125 s between groups. The participation of MNs in CMUGs and the order in which the segments are active during FWD and BWD crawling are inferred from the data (Figures 2 and 7).

Constraints are placed on the model parameters based on knowledge of the circuit. The nonzero elements of **J**^*p*^ and **J**^*m*^ are determined from the TEM reconstruction, and signs are constrained using neurotransmitter identity when available. If the neurotransmitter identity of a neuron is not known, a cost term is used to promote sign consistency (see below). Time constants **τ** are constrained to be between 50 ms and 1 s (these represent combined membrane and synaptic time constants).

At the beginning of optimization, the biases **b**^*p*^ and **b**^*m*^ are initialized equal to 0.1 and 0, respectively. Time constants **τ** are initialized to 200 ms. **I**^*FWD*^ and **I**^*BWD*^ are initialized uniformly between 0.05 and 0.15 for each neuron. To initialize **J**^*p*^ and **J**^*m*^, initial connection strengths are taken in proportion to synapse counts from the TEM reconstruction with a scaling factor of **±0.005** for excitatory/inhibitory connections. Connections within a model segment are taken from the TEM reconstruction of A1, while connections from A1 to A2 or A2 to A1 are taken from the corresponding cross-segmental reconstructions (and are thus likely less complete than the within-segmental connectivity). To focus only on strong and reliable connections, those that comprise less than 5% of the synapses received by a post-synaptic neuron are ignored.

The cost function that is optimized consists of a term *C*_*targ*_ that penalizes deviations of the MN activities from their targets and regularization terms to promote realistic solutions. The first term is given by 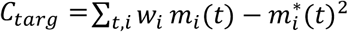, where 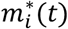 is the target activity for MN *i* and *w*_*i*_ is a weighting term, proportional to 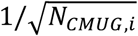 where *N*_*CMUG,i*_ is the number of neurons in the CMUG of neuron *i* (this scaling ensures the target patterns of CMUGs with few MNs are still reproduced accurately).

The regularization terms include a term *C*_*A*1*8b,A*2*7h*_ = 0.05 · (Σ_*t*∈*FWD*_ *p*_*A*18_ (*t*) + Σ_*t*∈*BWD*_ *p*_*A*27_ (*t*)), which suppresses the activity of the A18b and A27h neurons for behaviors during which they are known to be quiescent. The second regularization term *C*_*seg*_ constrains PMN activity to reflect the timing of segmental activation. It is equal to

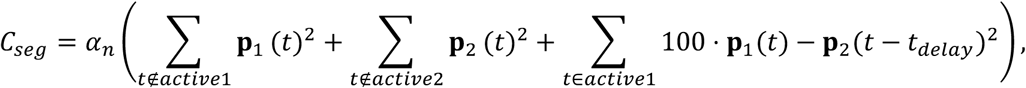

where active1 and active2 represent the times at which segments 1 and 2 are active, and *t*_*delay*_ is the time delay between segment 1 and 2 activations (equal to −1 s for forward waves and +1 s for backward waves). The last term in this equation ensures that PMN activity in the A1 and the A2 segments is similar but offset in time. The scaling term *α*_*n*_ increases quadratically from 0 to 0.01 over the 1000 training epochs. The final term *C*_*sign*_ = *α*_*n*_ Σ_*i,j,k*_[− *J*_*ik*_*J* _*jk*_]_+_ penalizes connections of inconsistent sign from neurons whose neurotransmitter identity is not known (**J** here represents all connections, including those onto MNs or PMNs).

The total cost, equal to the sum of the terms described above, is optimized using the RMSProp optimizer for 1000 epochs. During each epoch, the costs corresponding to one FWD and one BWD trial are averaged. The learning rate decreases from **10**^−2^ to **10**^−3^ logarithmically over the course of optimization.

## Supporting information

Supplemental Table 1. PMNs

Supplemental Table 2. Neurotransmitters

## Acknowledgements

We thank Luis Sullivan, Emily Sales, and Hiroshi Kohsaka for comments. B.M. was supported by an NIH training grant T32HD007348. A.C. was supported by HHMI. A.L.-K. was supported by the Burroughs Wellcome Foundation, the Gatsby Charitable Foundation, the Simons Collaboration on the Global Brain, and NSF award DBI-1707398. C.Q.D. and A.A.Z were supported by HHMI, where C.Q.D. is an Investigator.

---

**Figure 2 – Supplement 1a,b.**
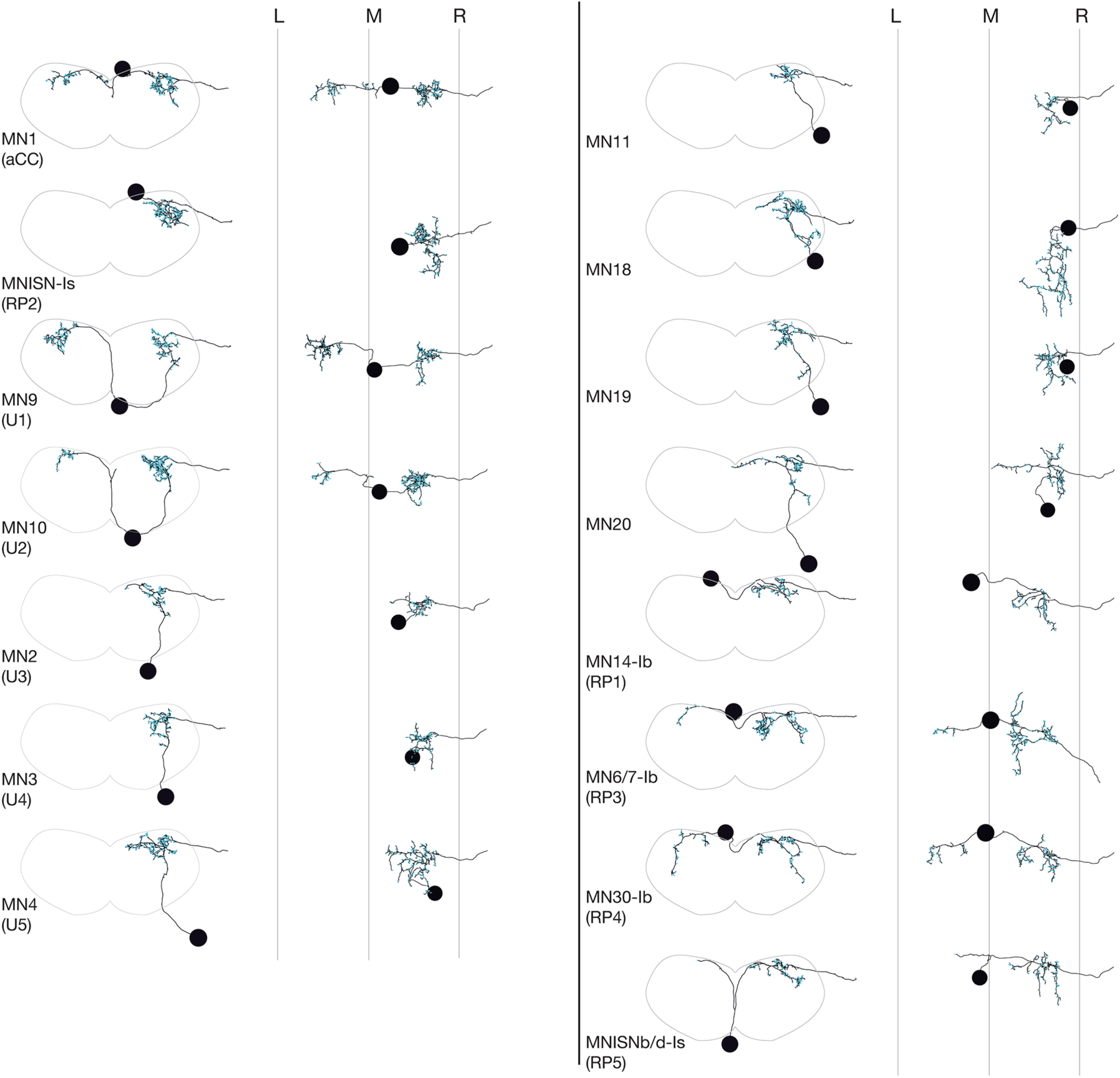
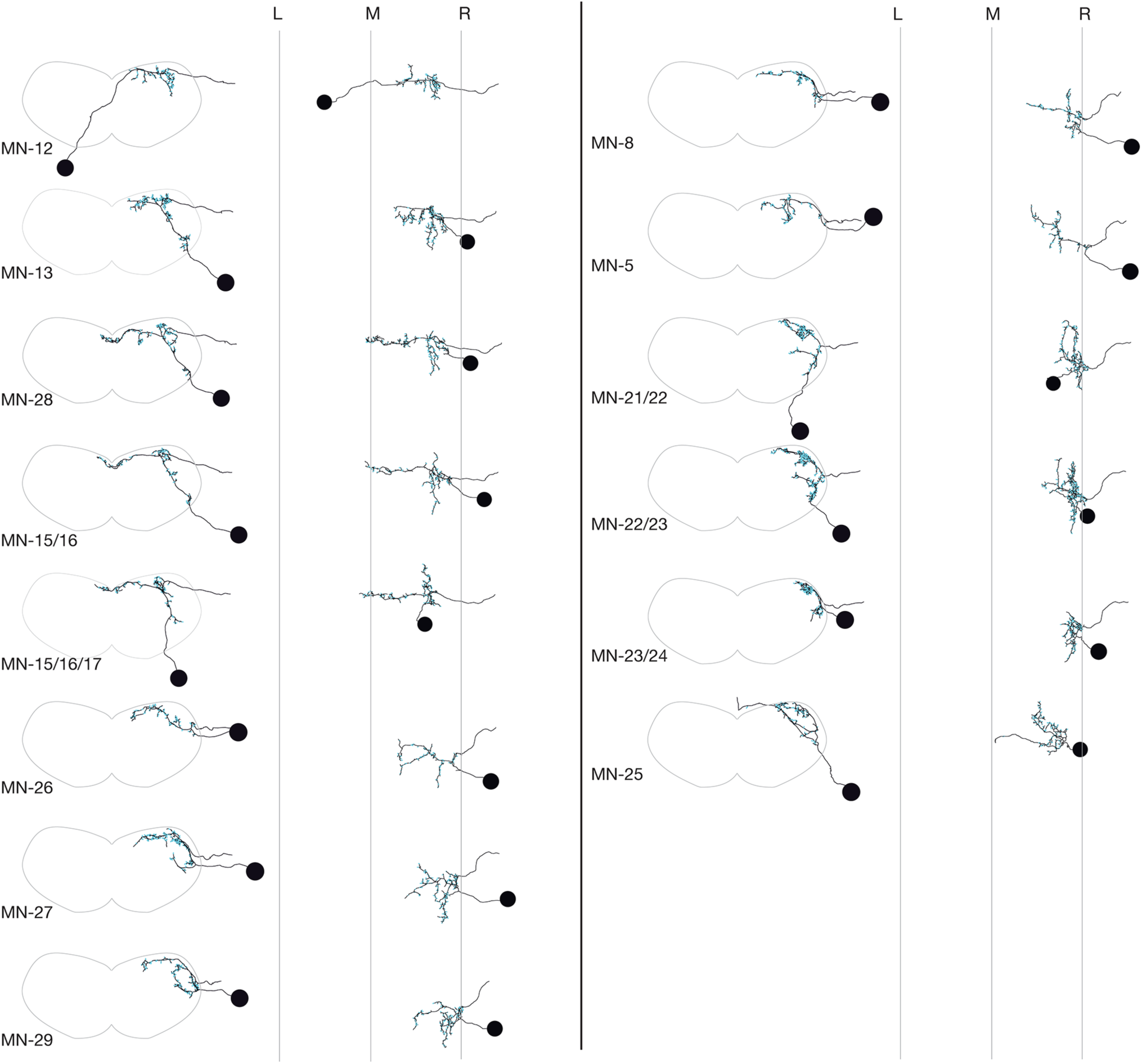
Reconstruction and Identification of A1 MNs using ssTEM. MN names are shown in bottom left for each reconstruction. The morphology of each MN is shown in cross-sectional (left) and dorsal view (right). In the cross-sectional views, neuropil boundary is shown in gray. In the dorsal views, L, M, R stand for Left neuropil border, Midline, Right neuropil border, respectively. Cyan dot depicts the post-synaptic sites on MNs.

**Figure 3 – supplement 1.**
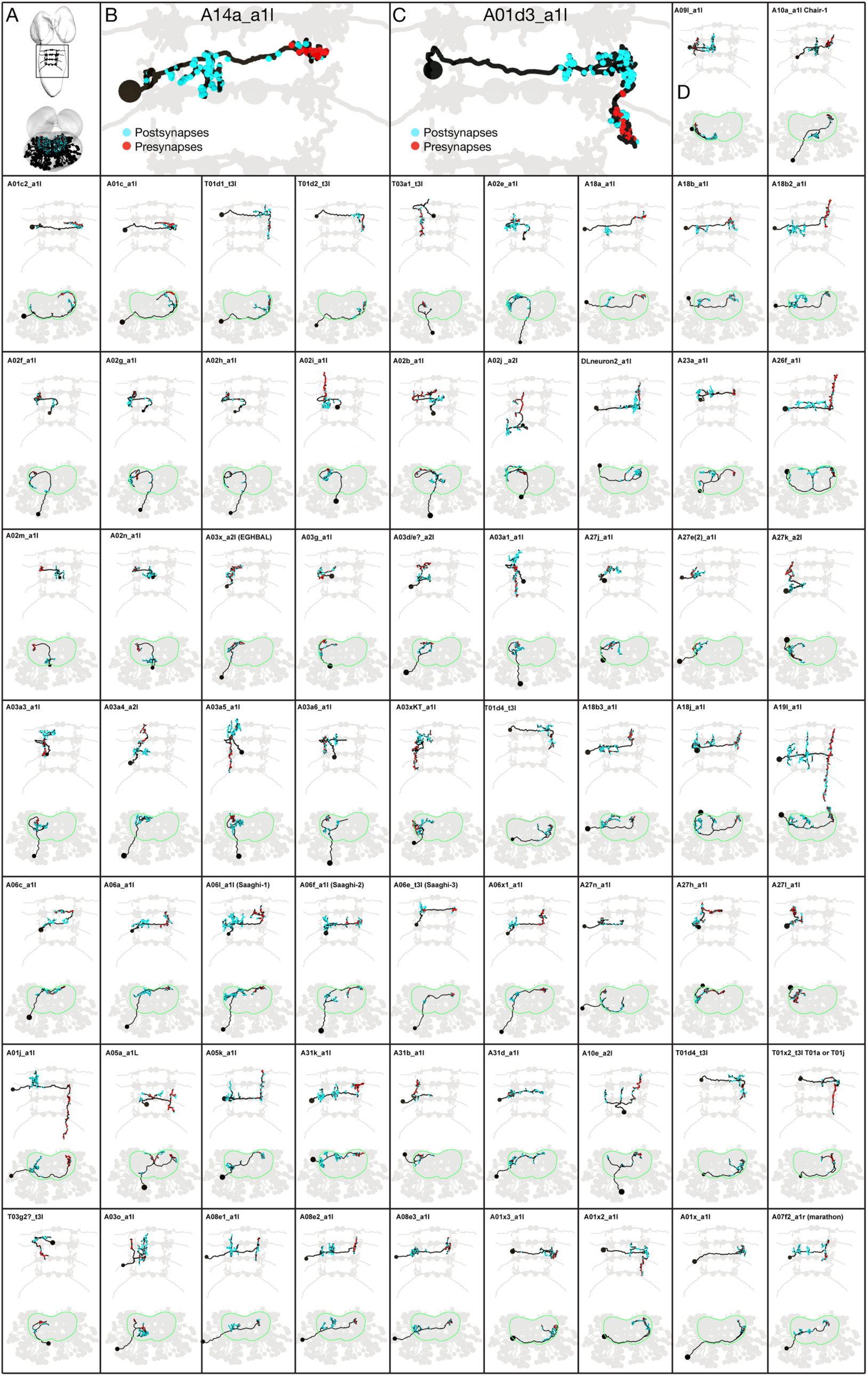
All premotor neurons traced in the TEM volume. (A) Dorsal view (top) and posterior view (bottom) of all 67 premotor neurons. (B-C) Magnified morphology of A14a (B) and A10d3 (C) PMNs. Red and cyan dots indicate pre-synaptic and post-synaptic sites respectively, and light-gray shaded neurons are MN1(aCC) in four subsequent segment (A1-A3) to define anteroposterior segment boundaries in the VNC. (B) In A14a PMN, most of post-synaptic sites are located in the proximal side (same side as the cell body), while pre-synaptic sites are located in the distal side (away from the cell body). (C) In A01d3 PMN, both of post-synaptic and pre-synaptic sites are located in the distal side (away from the cell body). (D) Individual PMNs. Names in upper left corner. In all panels, neuron names are at the top; red and cyan dots indicate pre-synaptic and post-synaptic sites respectively, and light-gray shaded neurons are MN1(aCC) in four subsequent segment (T3-A3) to define anteroposterior segment boundaries in the VNC.

**Figure 3 – supplement 2.**
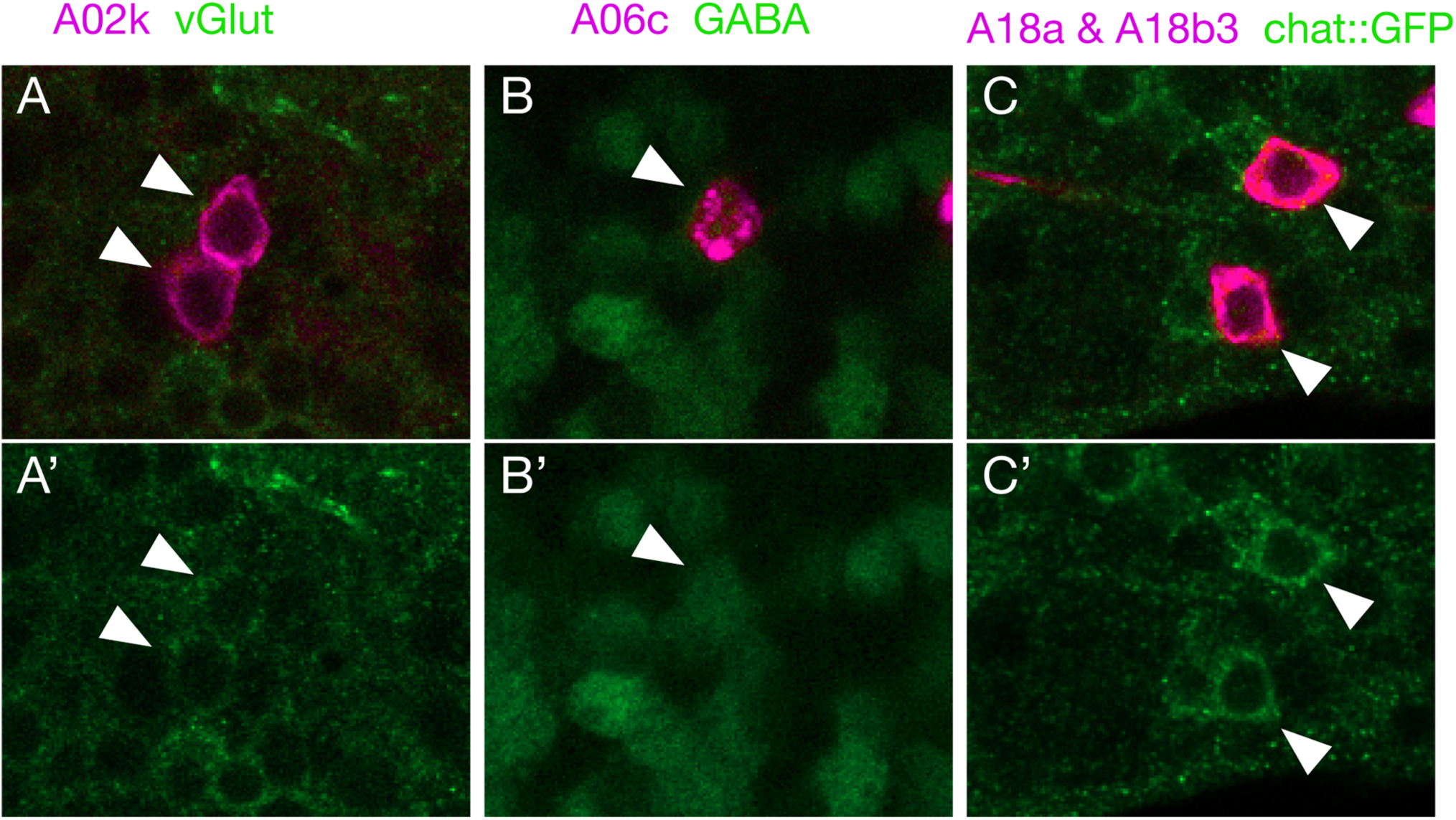
Neurotransmitter expression in premotor neurons. (A) A02k (magenta) is positive for vesicular glutamate transporter (vGlut) staining (white arrowheads) consistent with an inhibitory function. (B) A06c (magenta) is positive for GABA staining (white arrowheads), consistent with an inhibitory function. (C) A18a (top neuron) and A18b3 (bottom neuron) are positive for Chat:GFP staining (white arrowheads) indicating they are cholinergic, consistent with an excitatory function. See Table 3 for the neurotransmitter profile of all assayed neurons.

**Figure 5 – supplement 1.**
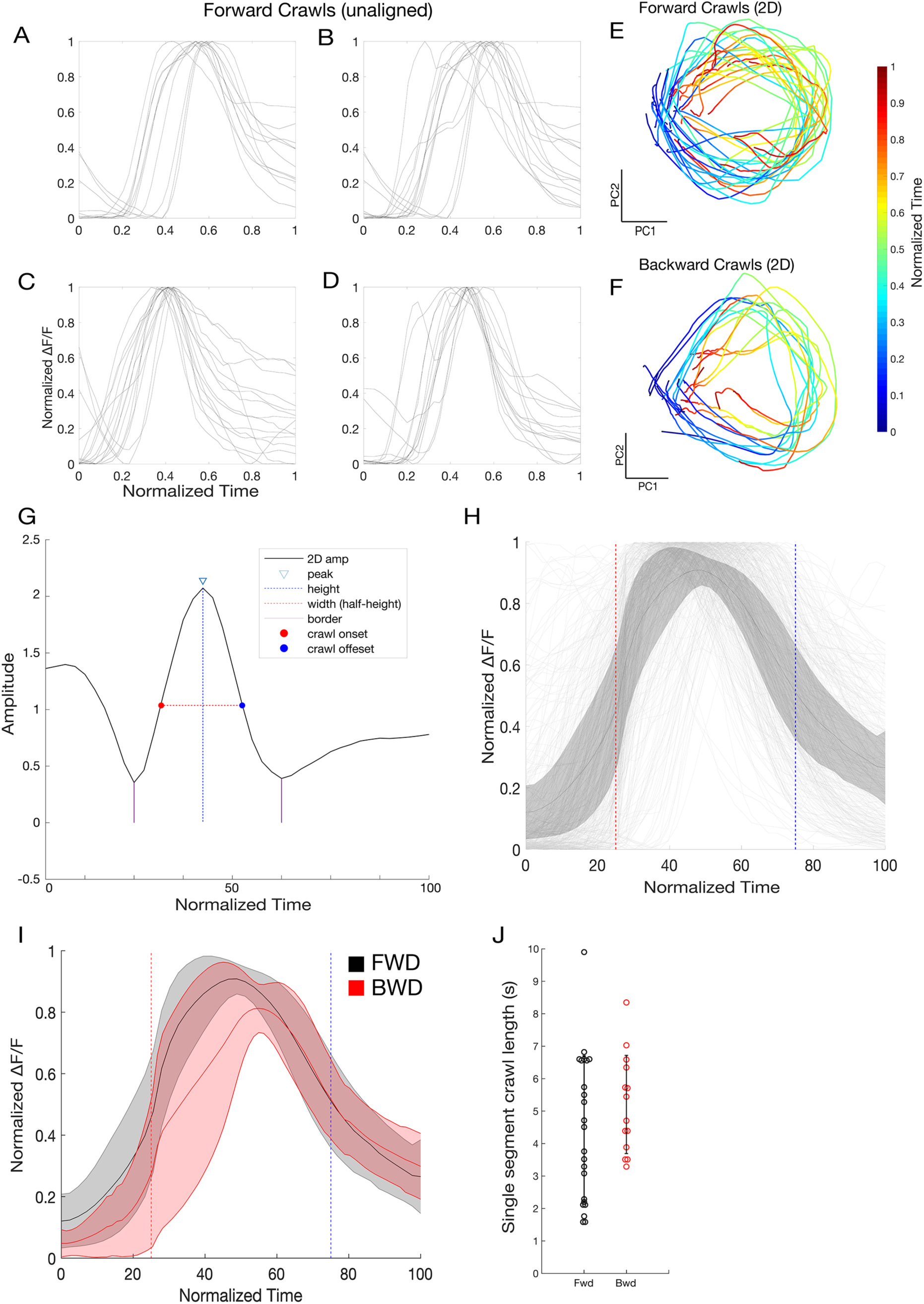
PCA-based alignment of crawl cycles. (A-D) Plots of four representative forward crawls show a high degree of variability in the crawl structure. (E-F) 2D projection of forward and backward crawl cycles. Crawl cycles are represented as rotations away from and back towards the origin. Color changes (from blue to red) represent time. Directionality is not uniform given that the group of analyzed muscles in each crawl cycle is different in each case (all crawls used had at least 40% of the muscles analyzed in the segment). (G) Amplitude of a representative forward crawl in the same 2D space. The peak of this activity was defined as the center of a crawl cycle, and the peak width at half the height of the peak was used to find crawl-start and crawl-end times. (H) All analyzed forward muscles aligned. Grey lines represent individual muscles, black line represents the average activity of all muscles with error bars representing standard deviation. Red dotted line represents the crawl-start alignment point, and the blue line represents the crawl-end alignment point. (I) Average activity of forward (black) and backward (red) crawls across all experiments. (J) Single segment crawl length determined for each crawl (n = 24 forward / 14 backward). Crawl length is determined by calculating the width of the 2D representation of the crawl cycle.

**Figure 5 - supplement 2.**
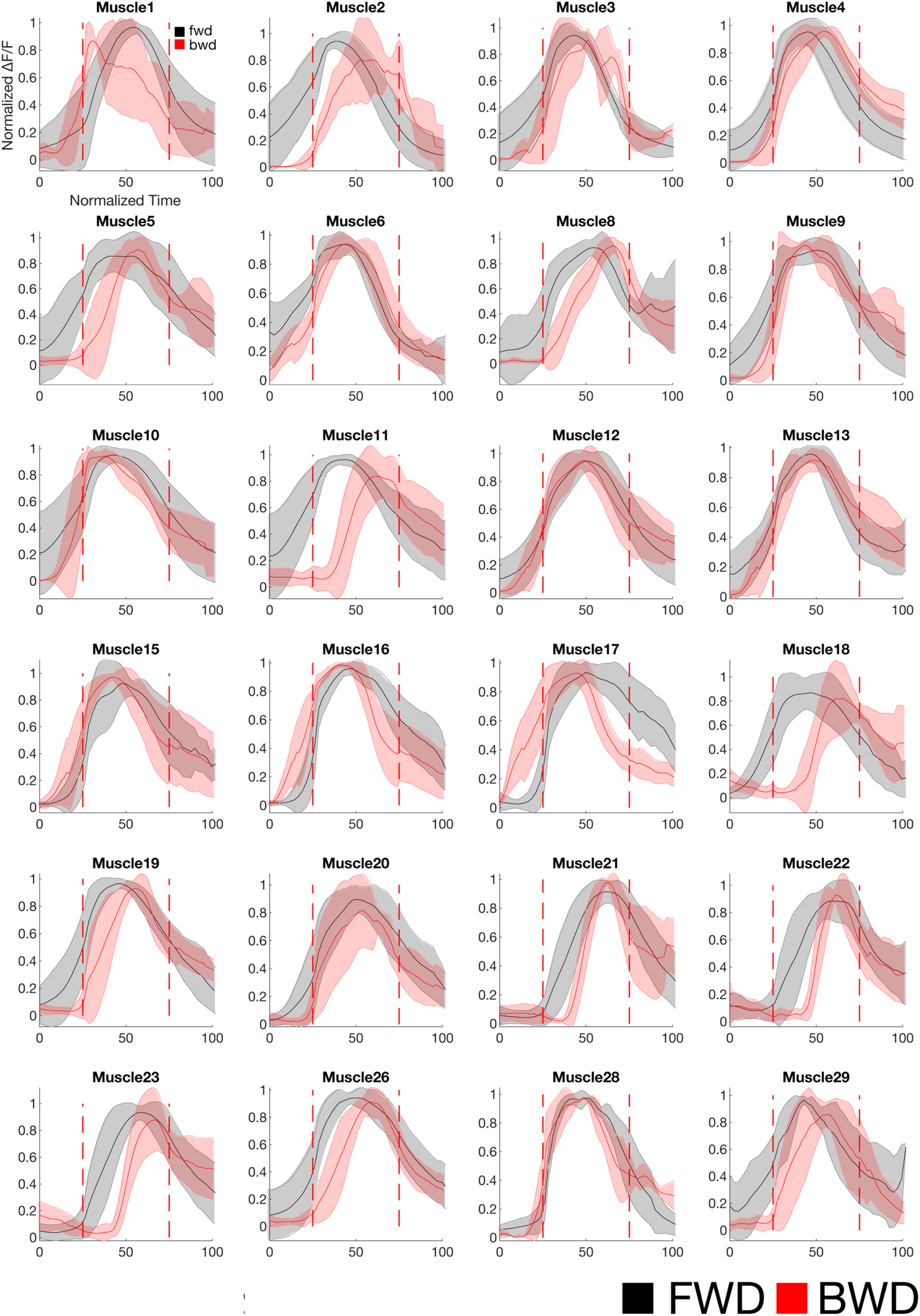
Average activity of all muscles analyzed during forward and backward locomotion. Red dotted lines represents the onset and the offset points. Red solid line represents the average activity of the muscles during backward with error bars (light red ribbon) representing the standard deviation. Black line represents the average activity of the muscles during forward with error bars (gray ribbon) representing the standard deviation. Some muscles show similar activity timing during both forward and backward locomotion (e.g. 10, 12, 13, 28), while some show differential activity timing during forward and backward locomotion (e.g. 1, 5, 18, 26, 29). Note that although muscle 21, 22, and 23 are activated as last CMuGs during both forward and backward locomotion, overlay of their activity in forward and backward locomotion indicates that these muscles are activated later in backward than forward. Therefore, the interval between CMuG4 and CMuG 1-3 is longer in backward than forward locomotion.

**Figure 10 – Supplement 1.**
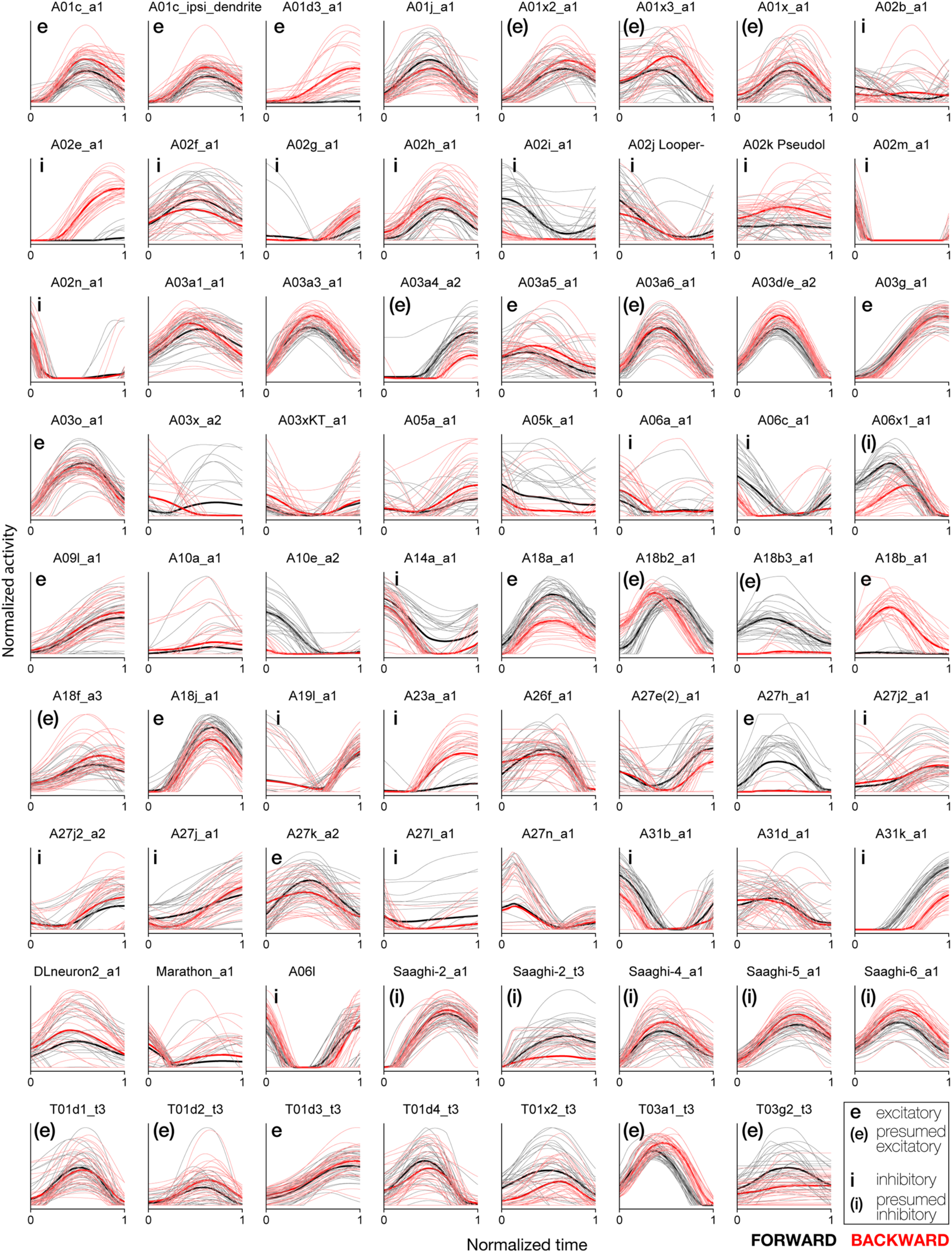
Recurrent network model of PMN activity during forward and backward locomotion. Normalized activity of PMNs in the model during forward and backward crawling. Thick lines denote averages over the ensemble of models generated. Time is measured relative to MN onset and offset as in Figure 10B. e, excitatory;(e), presumed excitatory based on lineage; i, inhibitory; (i), presumed inhibitory based on lineage.

**Supplemental File 1**. CATMAID .json file of all reconstructed motor neurons in segment A1 as of 17 February 2019.

**Supplemental File 2**. CATMAID .json file of all 67 pair of reconstructed pre-motor neurons in segment A1 as of 17 February 2019.

